# Evidence of convergent evolution in the nuclear and mitochondrial OXPHOS subunits across the deep lineages of Squamata

**DOI:** 10.1101/2024.11.14.623538

**Authors:** Oscar Wallnoefer, Alessandro Formaggioni, Federico Plazzi, Marco Passamonti

## Abstract

The OXidative PHosphorylation System (OXPHOS) is composed of subunits encoded by both the nuclear and mitochondrial genomes, which are subject to distinct evolutionary pressures. Nevertheless, the cooperation between OXPHOS subunits is essential for proper OXPHOS function, as incompatibilities between subunits can be highly deleterious.

The order Squamata is a good candidate for studying unusual patterns of mitochondrial evolution. The lineages leading to the snake and agamid clades likely experienced convergent evolution in mitochondrial OXPHOS genes, potentially linked to their distinctive feeding strategies. This deep signal of convergence can also be inferred from mitochondrial markers, which provide strong support for the monophyly of these two groups.

In the present study, we annotated the mitochondrial and nuclear OXPHOS genes of 56 Squamata species. The nuclear OXPHOS subunits that physically interact with mitochondrial proteins also support the clade clustering snakes and agamids. Additionally, we found a significant number of convergent amino acid changes between agamids and snakes, not only in mitochondrial OXPHOS genes but also in nuclear ones, with a higher rate of convergence in the nuclear OXPHOS subunits that play central roles in the OXPHOS complexes.

Overall, the common selective pressures in two distinct lineages can lead two sets of genes, encoded by two different genomes, to exhibit similar patterns of convergent evolution, affecting the phylogenetic signal of these genes. Thus, we highlight how the phylogenetic signal of OXPHOS genes, through the coevolution of subunits and their adaptation to specific evolutionary pressures, can be influenced and may diverge from the signal supported by most other genes.

**GRAPHICAL ABSTRACT:** In Squamata, nuclear genes support the monophyly of Pleurodonta and Acrodonta. However, OXPHOS genes, bot nuclear and mitochondrial, place Acrodonta in sister relationship with Serpentes. This phylogenetic discordance is likely due to convergent evolution along the that leading to Serpentes and Acrodonta.

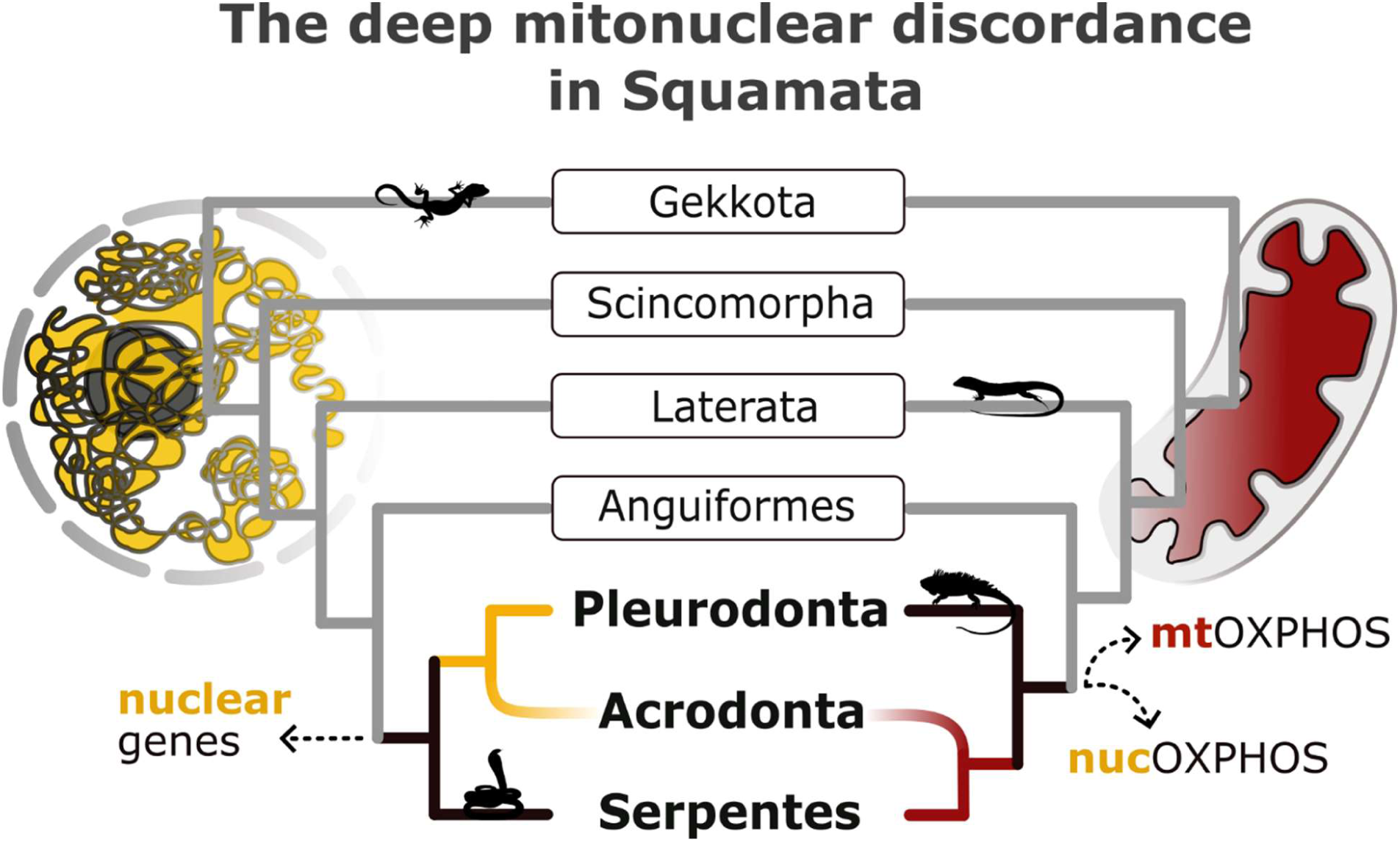

## 1.0 INTRODUCTION

The dual genomic nature of eukaryotes has intrigued evolutionary biologists for decades, raising fundamental questions about the coevolution of nuclear and mitochondrial genomes. Approximately two billion years ago, endosymbiosis between an archaeon and an alpha-proteobacterium led to the emergence of eukaryotic cells (Williams and Embley 2014). This event triggered a dynamic balance of cooperation and conflict between the two distinct genomes within the same cellular environment, resulting in the evolution of mitochondria as energy-producing organelles with their own DNA. While most genes from the alpha-proteobacterium were either lost or transferred to the nuclear genome, a conserved small set of mitochondrial genes was retained in the mitochondrial DNA (mtDNA), which varies across different eukaryotic lineages (Formaggioni et al. 2021). In most bilaterian animals, protein-coding genes are limited to 13 genes that encode essential subunits of the oxidative phosphorylation (OXPHOS) pathway (Hill 2019). The OXPHOS pathway is not exclusively determined by mitochondrial genes (mtOXPHOS) and the minimum set of subunits necessary to assure a well-functioning electron transport chain is never completely encoded in the mtDNA (Adams and Palmer 2003). In Metazoa, about 70 nuclear encoded OXPHOS genes (nucOXPHOS) also contribute to the five involved complexes. Four complexes constitute the electron transport system (CI, CII, CIII, CIV), while the fifth one is the ATP synthase (CV). Whereas CII is strictly nucleus-encoded, the other four complexes are assembled from mitochondrial and nuclear OXPHOS subunits. Each complex has a core, mainly made by mtOXPHOS subunits, which plays the catalytic activity (Hill 2019). Since most of the nucOXPHOS subunits mediate the stabilization and regulation of the complex, these subunits are called “supernumerary” (Signes and Fernandez-Vizarra 2018; Pfanner et al. 2019).

Literature postulates that the critical importance of the OXPHOS pathway, including tight coupling between catalytic centers and the precise interactions between subunits, leads to concerted evolution among OXPHOS genes (Sloan et al. 2018; Hill 2019). Nuclear compensation is the best described mechanism, though not entirely demonstrated (Hill 2019), among the plethora of various processes driving mito-nuclear coevolution (Nabholz et al. 2013; Havird and Sloan 2016; Hill 2019; Wernick et al. 2019). This hypothesis posits that selection on counterbalancing mutations plays a significant role in sustaining the OXPHOS functionality, which suffers the higher evolutionary rate of the mitochondrial genes in bilaterians (Lynch 2010; Popadin et al. 2013).

One of the most persuasive lines of evidence derives from a study on primates by Osada and Akashi (2012), which demonstrated that rapid evolution in some nucOXPHOS genes reflects adaptive responses to counteract slightly deleterious mtOXPHOS substitutions. However, empirical evidence regarding the role of compensatory mutations in nucOXPHOS genes remains limited to few studies and contradicted by others (Zhang et al. 2013; Piccinini et al. 2021; Weaver et al. 2022).

Coevolution and interaction of proteins are topics of great interest among evolutionary biologists, and many approaches were developed to investigate them (de Juan et al. 2013). Among these, the Evolutionary Rate Correlation (ERC) is one of the most used approaches and has provided several pieces of evidence supporting the coevolution of proteins that are part of the same metabolic pathway or that participate in the same cellular functions (Clark et al. 2012; Clark et al. 2013; Steenwyk et al. 2020). For instance, higher substitution rates occur in mitochondria-associated nucleus-encoded proteins in lineages with faster mitochondrial evolution and tend to show high covariations (Havird and Sloan 2016). ERC was already investigated on OXPHOS genes in three large clades among Metazoa: insects (Yan et al. 2019), bivalves (Piccinini et al. 2021) and mammals (Weaver et al. 2022). All of them registered similar evolutionary rates between OXPHOS proteins in comparison to nuclear genes that are not associated with mitochondria. In particular, contact nucOXPHOS seem prone to support stronger correlation with the amino acid substitution rate of mtOXPHOS, according to the hypothesis that physical interactions contribute to ERC as the effect of maintenance of proper binding sites (Goh et al. 2000; Ramani and Marcotte 2003; Salmanian et al. 2020).

Here, we investigated mitonuclear coevolution in Squamata, a clade selected for its distinctive features. Squamata is one of the most diverse orders among chordates, with over 12,000 living species (Uetz et al. 2023), and has undergone a highly successful radiation since Jurassic, surviving to the end-Cretaceous mass extinction (Evans and Jones 2010). This clade is notably controversial from a phylogenetic perspective, often displaying extreme phenotypic variability that contrasts with molecular data, complicating efforts to reconcile historical morphological hypotheses with molecular phylogenetics (Burbrink et al. 2020; Singhal et al. 2021).

Additionally, mitochondrial markers can sometimes contradict nuclear ones, leading to mitonuclear discordances (Albert et al. 2009). A notable example of deep mitonuclear discordance involves the relationship between Acrodonta, a clade that includes chameleons and agamids, Pleurodonta, which includes iguanas, and Serpentes (snakes). Analyses based on mitochondrial markers have repeatedly placed Acrodonta in a sister relationship with Serpentes (Böhme et al. 2007; Albert et al. 2009; Castoe et al. 2009). In contrast, analyses based on nuclear markers support a closer relationship between Acrodonta and Pleurodonta, forming a clade known as Iguania (Burbrink et al. 2020; Singhal et al. 2021).

Mitonuclear phylogenetic discordances are quite common among Metazoa and can be attributed to differences in inheritance mechanisms, recombination rates, mutation rates, effective population sizes, and evolutionary constraints between the two genomes (Toews and Brelsford 2012). The organellar environment may exert radically different pressures on mtDNA compared to nuclear DNA, potentially leading the two genomes along divergent evolutionary paths. Incoherent topologies may also arise from factors such as mutational constraints, asymmetric selection, introgression, hybridization, environmental pressures, or demographic peculiarities, such as sex-biased dispersal (Funk and Omland 2003; Toews and Brelsford 2012). However, while many instances of mitonuclear discordance are documented at the species or genus level, the deep node discordance within the Squamata order is unique and particularly intriguing. According to Castoe and colleagues (2009), an ancient event of adaptive convergent evolution on the OXPHOS may explain the monophyly of agamid lizards and snakes. This hypothesis is corroborated by signals of positive selection in the mitochondrial OXPHOS genes in snakes (Castoe et al. 2008). This selective pressure may be due to metabolic adaptations related to the unique feeding habits of snakes, which consume infrequent, but large meals that result in fluctuations in oxygen consumption of up to 4,000% (Secor and Diamond 1998). The authors suggest that the convergence between snakes and agamid lizards might be due to similar pressures on mtOXPHOS and hypothesize the existence of similar signals on other genes involved in the same pathway (Castoe et al. 2009).

We selected 56 squamate species belonging to 22 different families to investigate several key aspects of mitonuclear coevolution. First, we tested the phylogenetic signal among genes involved in the OXPHOS by performing phylogenetic analyses with both mitochondrial and nuclear OXPHOS genes. As also evidenced in bivalves, phylogenies based on different sets of OXPHOS genes can produce mirrored topologies, supporting the hypothesis of mitonuclear coevolution (Formaggioni et al. 2022). Second, we examined the covariation in evolutionary rates between nuclear OXPHOS, mitochondrial OXPHOS and a set of nuclear control genes. Finally, given the pronounced convergence signal between agamid lizards and snakes on mtOXPHOS genes, we searched for molecular evidence of similar convergence in nucOXPHOS genes, hypothesizing that strong evolutionary pressure on mtOXPHOS would also affect interacting nuclear subunits. Overall, the clade Agamidae+Serpentes is strongly supported by mitochondrial and by the subset of nuclear OXPHOS subunits that closely interact with mitochondrial counterparts. Some OXPHOS subunits show clear signatures of convergent evolution. This suggests that the OXPHOS system has been under similar selective pressure in both branches.

## 2.0 RESULTS

### 2.1 The OXPHOS datasets

The OXPHOS genes were annotated on all the Squamata genome assemblies available on NCBI at March 2023, limiting the number of congeneric species. 10 species were excluded because we failed to assemble the mitochondrial genome. Additionally, 3 species were excluded because BUSCO completeness scores were below 80%. In total, the dataset comprehends 56 Squamata species plus three reptilomorph outgroups, namely Sphenodon punctatus (Rhynchocephalia), Alligator mississippiensis (Crocodilia) and Caretta caretta (Testudines) (see Table S1). The completeness matrix of mtOXPHOS genes (Fig. S2a) was very high, with only 7 genes missing out of a total of 767 possible genes (13 genes times 59 species). Regarding nucOXPHOS genes, 3,778 (87.7%) out of the expected 4,307 genes (73 genes times 59 species) were successfully annotated (Fig. S2b). Namely, some subunits from the KEGG list (map00190) were not found or were misannotated; consequently, a total of 73 nuclear subunits was included in the analysis. A third matrix included all the single-copy genes inferred with BUSCO using the Metazoa_odb10 dataset, filtering out genes that were absent in more than 20% of the OTUs. The final dataset is comprised by 896 BUSCO genes and will henceforth be called the BUSCO genes dataset.

### 2.2 Interacting OXPHOS genes share the same phylogenetic signal

We inferred the phylogenetic signal from mtOXPHOS makers using the Bayesian and the Maximum Likelihood (ML) approaches on the nucleotide and amino acid datasets. The Bayesian approach on the nucleotide dataset almost confirmed the mitochondrial topology proposed by Castoe and colleagues (2009). The only difference is that in our analysis Anguiformes is the first clade to separate from all the other Toxicofera (snakes, iguanians, and anguiforms), although with a non-significant posterior probability (PP) of 0.91, rather than being in sister relationship with Pleurodonta. Many differences were found between topologies inferred using different methods (i.e., ML) or character types (i.e., nucleotides or amino acids). The four topologies differed particularly in the deep relationships and many deep nodes received low support values (Fig. S3c- f). Teiidae and Lacertidae were inferred as sister groups only in the Bayesian analysis using the nucleotide dataset, while the ML analyses placed Teiidae in sister relationship with Agamidae+Serpentes. Gekkota and Scincomorpha were inferred in a basal position only in the Bayesian analyses, while in the ML trees their position is highly variable, especially for Scincomorpha. We performed tests to exclude that some markers were oversaturated by multiple substitutions, especially in the nucleotide datasets. However, we did not detect notable differences on the third base in the number of substitutions between an uncorrected model and the Tamura model, which accounts for multiple substitutions (Fig. S4). Although the four topologies inferred from the mtOXPHOS largely differed in the resolution of some nodes, the clade Agamidae+Serpentes has been inferred in all four trees, with high support values.

Regarding the nucOXPHOS dataset, the ML analysis on nucleotides identified Agamidae as the sister group of Pleurodonta, while amino acids support the mitochondrial topology (Fig. S3g, h). The phylogenetic position of Lacertidae and Teiidae was ambiguous: using nucleotide data, these families formed a clade that was the sister group of Toxicofera (Fig. 1a, S3g), whereas, using amino acid data, Agamidae and Teiidae were close to snakes (Fig. S3h). We then narrowed the dataset to the nucOXPHOS subunits that are physically in contact with the mitochondrial counterparts (contact nucOXPHOS). The ML tree inferred from this dataset placed Agamidae in a sister relationship with Serpentes, in accordance with the mtOXPHOS dataset (Fig. 1b, S3i).

**Figure 1.**
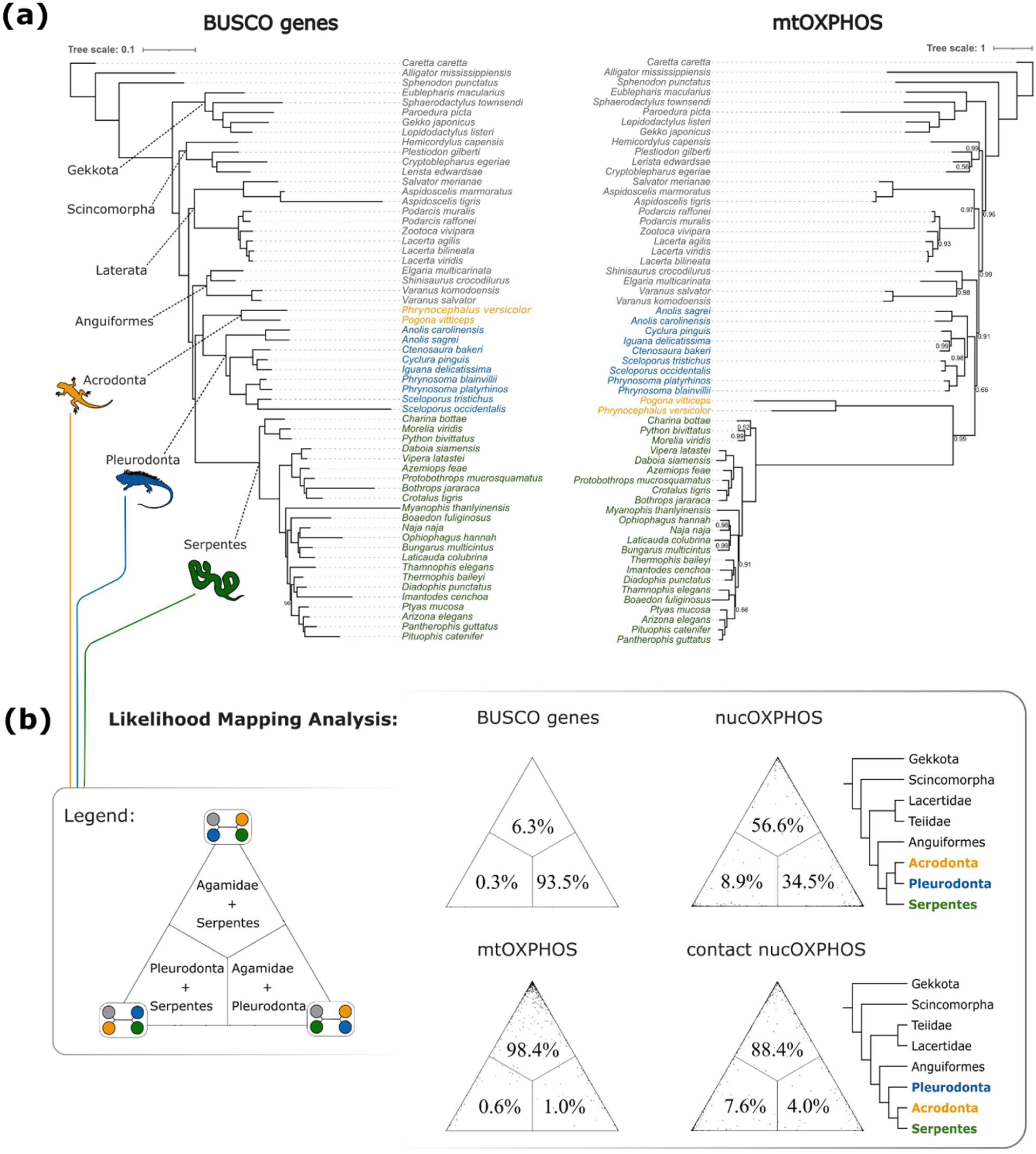
The phylogenetic analysis of mitochondrial and nuclear markers. a) The mitonuclear discordance affects the Agamidae (Acrodonta) position. The monophyletic origin of Iguania (Acrodonta+Pleurodonta) is supported by BUSCO genes (left, ML tree), but rejected using mitochondrial OXPHOS genes (right, Bayesian tree). The formers support the sister relationship Acrodonta+Serpentes. Only poster probabilities under 100 are reported in the tree. b) The placement of Agamidae within the Squamata phylogeny varies incoherently when analyzing BUSCO genes, mtOXPHOS genes, nucOXPHOS genes, and a subset of nucOXPHOS genes whose subunits closely interact with mitochondrial counterparts (referred to as contact nucOXPHOS). The Likelihood Mapping Analysis evaluates the phylogenetic support for three alternative topologies: Agamidae+Serpentes (top angle), Agamidae+Pleurodonta (bottom-right angle), and Pleurodonta+Serpentes (bottom-left angle). Schematic representations of the nucOXPHOS and contact nucOXPHOS topologies are shown.

In order to test the phylogenetic signal associated to each set of genes, we divided the OTUs into four groups (i.e., Agamidae, Serpentes, Pleurodonta and others) and performed the Likelihood Mapping Analysis (LMA). In the BUSCO dataset, 93.5% of quartets support the Iguania clade (Agamidae+Pleurodonta; Fig. 1b). In the mtOXPHOS dataset, almost all quartets (98.4%) of OTUs randomly selected each from the four groups were highly informative in supporting the Agamidae+Serpentes hypothesis (Fig. 1b). On the other hand, while Agamidae+Serpentes received only 56.6% support from the LMA on nucOXPHOS, this percentage rose to 88.4% when the analysis was restricted to the contact nuclear subunits (Fig. 1b). Therefore, mitochondrial and contact nuclear markers resolve the relationships between Agamidae, Serpentes and Pleurodonta concordantly.

Overall, nucOXPHOS markers support two alternative topologies (i.e., Agamidae+Serpentes or Agamidae+Pleurodonta).

We calculated the Site-Specific Likelihood Support (SSLS) for both topologies (Fig 2a, b). Then, we subtracted the SSLS calculated for the Agamidae+Pleurodonta topology from the SSLS calculated for the Agamidae+Serpentes topology (ΔSSLS). Positive values of ΔSSLS indicate stronger support for Agamidae+Serpentes, with values above 0.5 indicating significative support (Castoe et al. 2009). Conversely, negative values of ΔSSLS indicate stronger support for Agamidae+Serpentes, with values below -0.5 indicating significative support. The support for Agamidae+Serpentes is mostly concentrated in the contact subunit sites, that generally show a more positive ΔSSLS compared to non-contact genes with medians of –0.537 and –0.734 (Mann-Whitney; p-value = 0.000357; Fig. 2c) for contact nucOXPHOS and non-contact nucOXPHOS, respectively (Fig. 2a, b; Fig. S5). The number of sites that strongly support the Agamidae+Serpentes hypothesis is 477 compared to the 693 sites that support the species tree topology. However, as shown in Figure 2c, support for the former is higher among contact genes (280 out of 477), while support for the latter is higher among non-contact genes (387 out of 693). The phylogenetic signal is more retained in the first and second codon positions than in the third one (Fig. 2b; Fig. S5a, b). If we consider sites with a strong support for the Agamidae+Serpentes, the first and second codon positions show an average ΔSSLS significantly higher than the average ΔSSLS of third position sites (Mann-Whitney test; p-value = 0.0065; Fig. S5a, b).

**Figure 2.**
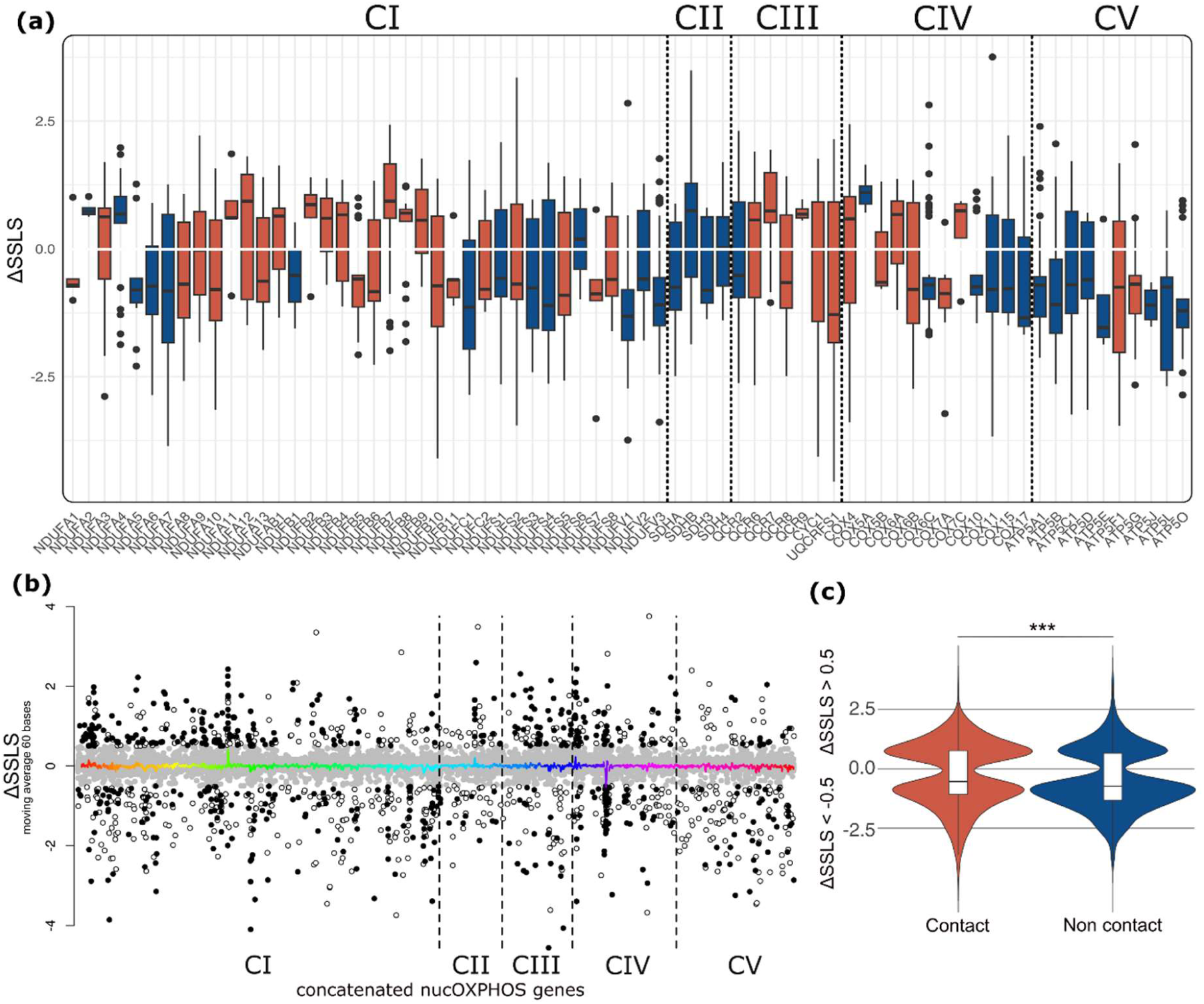
Site-wise support for the two discordant topologies. **a)** Boxplots of ΔSSLS restricted to informative sites (ΔSSLS > 0.5 and ΔSSLS < –0.5), associated to each nucOXPHOS genes and divided by complex. Positive values of ΔSSLS indicate support for the mitochondrial topology (Agamidae+Serpentes), while negative values support the monophyly of the Iguania clade. ΔSSLS > 0.5 and ΔSSLS <−0.5 are considered highly informative and constitute clues of strong support. Red and blue boxplots represent contact and non-contact nucOXPHOS, respectively. **b)** ΔSSLS subdivided into the first two codon positions (black) and the third one (white) associated to the concatenated nucOXPHOS dataset. Grey color indicates non informative sites. The moving averages (window size of 60 bases) are drawn with different colors. c) Violin plot of ΔSSLS divided by contact (red) and non-contact (blue) nucOXPHOS. The ΔSSLS associated to contact nucOXPHOS are significantly higher than the remaining nucOXPHOS.

### 2.3 Evolutionary rate correlations reveal signals of coevolution between mtOXPHOS and nucOXPHOS

We compared the evolutionary rates of the three datasets (i.e., mtOXPHOS, nucOXPHOS and BUSCO genes) across the 56 Squamata OTUs. Both OXPHOS datasets showed lower evolutionary rates than BUSCO genes in nearly all non-snake species, with mtOXPHOS genes being generally more conservative than the nucOXPHOS genes. Conversely, in Serpentes, the evolutionary rates of BUSCO genes were lower than those of OXPHOS genes, with the rates of the nucOXPHOS being generally lower than those of the mtOXPHOS (Fig. 3a). The same pattern was also observed in the agamid Phrynocephalus versicolor and, partially, in the agamid Pogona vitticeps. The similarity in evolutionary rates becomes even more evident when considering only the contact nucOXPHOS (Fig. 3a). Evolutionary Rate Correlations (ERC) are used to test whether genes have undergone similar evolutionary trajectories and functional relationships among proteins (de Juan et al. 2013). To ensure that the composition of the BUSCO dataset did not bias the analysis and to minimize the distorting effect of the dataset size, we normalized the root-to-tip distances of tress optimized on the mtOXPHOS, nucOXPHOS and BUSCO alignments using equal-sized random BUSCO subsets, as detailed in Methods. Then, we correlated the normalized mtOXPHOS, nucOXPHOS and BUSCO distances to each other. After computing 1,000 iterations, ERC between mtOXPHOS and nucOXPHOS reported a median Spearman’s ρ equal to 0.72 (Fig. 3b). Narrowing the nucOXPHOS dataset to the contact nucOXPHOS dataset, the median value of the ρ distribution significantly increased to 0.79 (Mann-Whitney test; p-value < 2.2×10^-16^). On the other hand, the ERCs calculated between BUSCO genes and mtOXPHOS and nucOXPHOS genes were 0.12 and 0.10 (median ρ), respectively.

**Figure 3.**
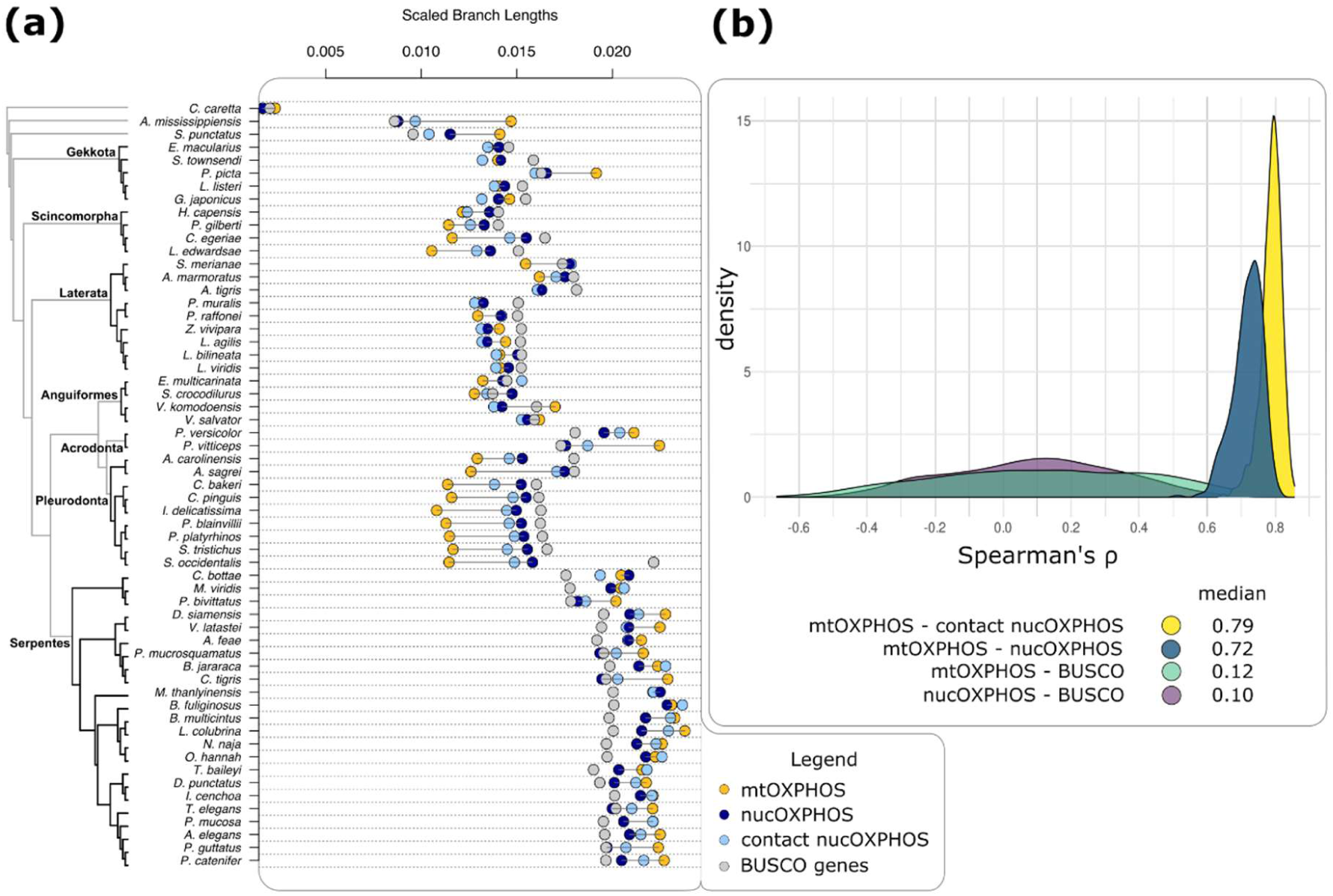
Evolutionary rates analyses. a) Comparisons between root-to-tip scaled distances of the four gene sets. mtOXPHOS and nucOXPHOS points are lined with a slight black line to highlight the distance between them. b) Density curves of Spearman’s ρ resulting from evolutionary rate correlations (ERCs) between nuclear and mitochondrial OXPHOS genes (see Materials). Distributions of Spearman’s ρ between OXPHOS genes and BUSCO genes were used as controls.

### 2.4 Signals of convergence

Since two independent clades, Agamidae and Serpentes, show similar patterns for both mtOXPHOS and nucOXPHOS genes compared to the BUSCO genes, we searched for signs of amino acid convergence between these two groups. We reconstructed all the non-synonymous substitutions along the Squamata phylogeny and compared all the possible pairs of independent branches. For each pair, we counted the number of convergent and divergent substitutions. We considered substitutions on both branches in the same codon, and we defined them convergent when the mutations led to the same amino acid, while we defined them divergent when they led to different amino acids. We also evaluated the rate of convergent substitutions relative to the total number of substitutions for each nucOXPHOS and mtOXPHOS gene.

We determined the linear relationship between the number of convergent substitutions and the total number of substitutions (i.e., convergent plus divergent substitutions along two lineages) for each gene, considering all possible pairs of independent branches. For each pair of branches, we calculated the difference between the observed number of divergences and the number of divergences predicted by the linear model (i.e., the residual of each point). We then estimated the distribution of these residuals and looked for the residual associated to the pair made by the branch leading to the Agamidae clade and the one leading to the Serpentes one (Fig 4a, b, c): this residual was often among the largest 2.5% residuals. Conversely, residuals associated to branches leading to either Agamidae or Serpentes and branches leading to one of the other major groups (namely, Gekkota, Scincomorpha, Laterata, Anguiformes, Pleurodonta) are rarely among the largest 2.5% residuals, and there is no consistent pattern involving a single pair of branches (Fig. S7a, b). This indicates a higher rate of convergent substitutions between Agamidae and Serpentes, which was not observed elsewhere. The same analysis on nucOXPHOS showed a similar pattern: 25 out of 73 subunits reported a significantly large residual involving Agamidae and Serpentes (Figure 4b). In particular, they were sixteen subunits of CI, four subunits of CIII, four subunits of CIV and one subunit of CV, while CII had no subunits showing this pattern. Even in this case, residuals associated to other deep branches were rarely included among the largest 2.5%.

**Figure 4.**
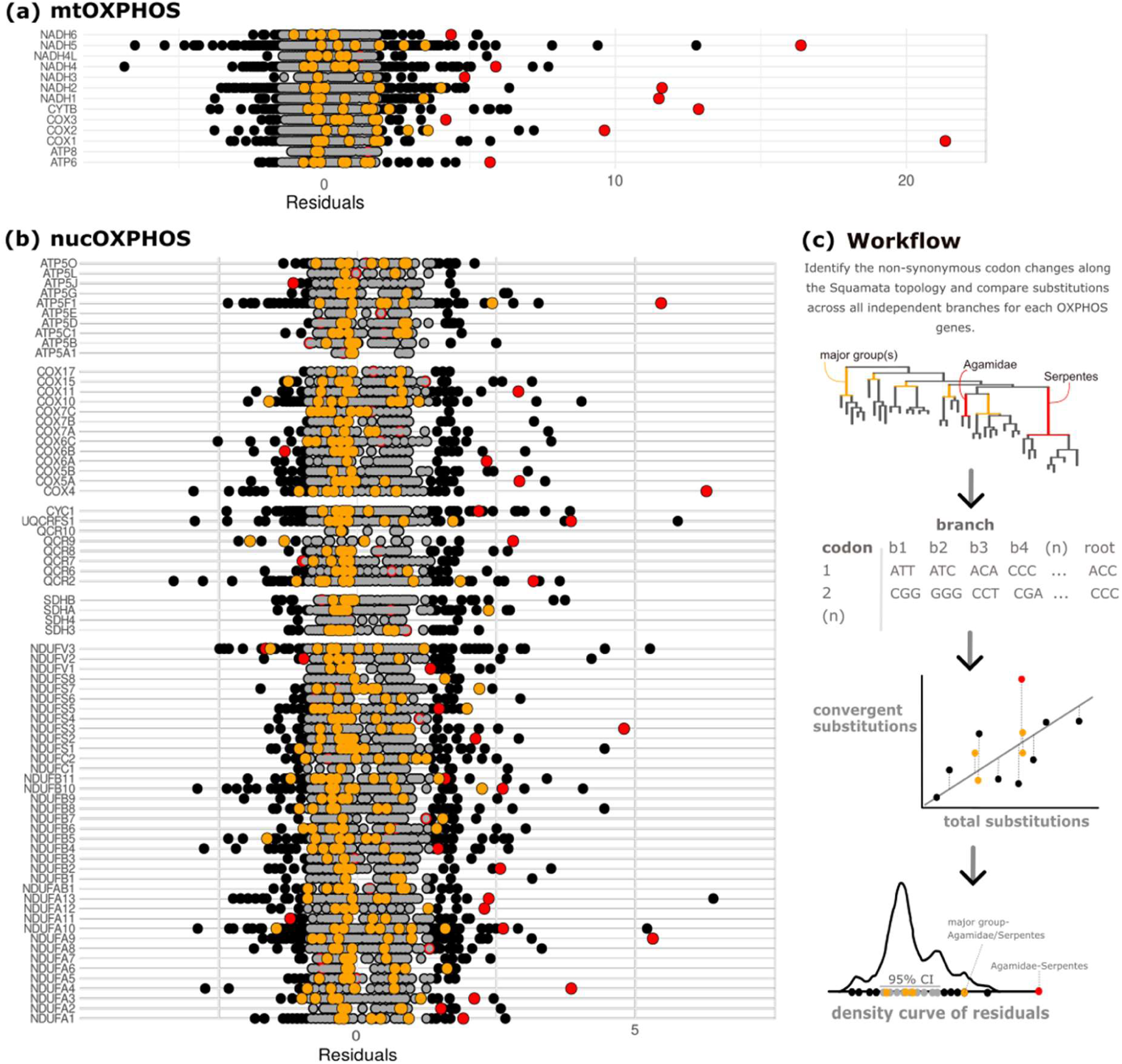
Convergent substitutions along the Squamata and Agamidae branches. Distributions of residuals associated to (a) the mtOXPHOS and (b) the nucOXPHOS genes. c) Schematic overview of the previous steps. We define “residual” the distance between the observed number of convergences out of total amino acid change and their expected ratio. We calculated the residuals distribution along the Squamata phylogeny for each codon of each OXPHOS genes. Points within the 95% confidence interval are grey. The pairs made by species of Agamidae and species of Serpentes are red. Control pairs are yellow.

## 3.0 DISCUSSION

Squamate reptiles are a good model to study mitonuclear coevolution: the deep phylogenetic discordance between mitochondrial and nuclear genes allows to test the strength of the phylogenetic signal applied to genes involved in the same metabolic pathway and that interact with each other. Moreover, it is possible to test how the adaptation measured on mitochondrial OXPHOS genes affects the nuclear counterparts. The phylogenetic trees inferred from our three Squamata datasets (i.e., mitochondrial OXPHOS genes, nuclear OXPHOS genes and BUSCO genes) were different, especially in the relationships between main branches.

Each matrix was analyzed with the ML and Bayesian inference methods using nucleotides or amino acids. Previous analyses showed that the inference method extensively influenced the topology from mtOXPHOS genes (Albert et al. 2009). However, both ML and Bayesian approaches confirmed the monophyly of Serpentes+Agamidae, confirming previous analyses on mitochondrial markers (Böhme et al. 2007; Albert et al. 2009; Castoe et al. 2009). Trees inferred from the nucOXPHOS dataset supported the monophyletic origin of Iguania using nucleotides (Fig. 1b). However, narrowing the analysis to the subset of genes that show interactions with mitochondrial subunits, we obtained a topology where Agamidae are sister to Serpentes (Fig. 1b, S7i), in accordance with the topology inferred from the mtOXPHOS dataset. This result indicates that the phylogenetic signal of interacting OXPHOS subunits converges towards the same topology.

A tight coevolution between nuc and mtOXPHOS subunits has also been confirmed by the ERC analysis (Fig. 3b). mtOXPHOS and nucOXPHOS datasets were significantly more correlated than the correlations between mtOXPHOS-random BUSCO genes and nucOXPHOS-random BUSCO genes, regardless the dataset used for normalization. Concordantly with the phylogenetic analysis, the subset of nucOXPHOS genes directly in contact with mitochondrial subunits showed the highest correlation with the mtOXPHOS dataset. Thus, even in Squamata, the coevolving subunits show higher ERC values, as previously demonstrated in insects (Yan et al. 2019), bivalves (Piccinini et al. 2021) and mammals (Weaver et al. 2022). Moreover, as in bivalves (Formaggioni et al. 2022), the phylogenetic discordance observed in mitochondrial markers has also been identified in some OXPHOS nuclear markers.

The necessity of maintaining physicochemical complementarity between interacting residues may influence the phylogenetic signal and evolutionary rates of subunits (Marmier et al. 2019). However, mirrored trees and correlated ERC might also be driven by global shared pressures arising from shared evolutionary histories, and not only from the presence of contact residues, (Goh et al. 2000; Lovell and Robertson 2010; Qin and Colwell 2018; Little et al. 2024). In this context, nucOXPHOS subunits would share the mtOXPHOS topology not because they are in contact with their mitochondrial counterparts, but because they are part of the complex structural core, unlike others that are more peripheral and farther from the catalytic centers (Signes and Fernandez-Vizarra 2018). Additionally, because the interface of interacting proteins typically makes up only about 10% of residues, and this proportion further decreases when focusing on binding sites (Lovell and Robertson 2010), the substantial coevolution detected using contact sequences might be obscured by the broader variability across entire domains/genes (Little et al. 2024). Notably, these two hypotheses are not mutually exclusive, and both could contribute to the observed phylogenetic signal.

Phylogenetic analyses, ERC and the surprisingly high rates of amino acid convergences confirmed the tight coevolution between mitochondrial and nuclear OXPHOS subunits, but why do some OXPHOS subunits support the clade Agamidae+Serpentes? Castoe and colleagues (2009) suggested that both lineages experienced similar selective pressures on OXPHOS genes, likely related to the peculiar feeding habits of these animals. Interestingly, in most Serpentes and Agamidae species, the evolutionary rates of OXPHOS genes, both nuclear and mitochondrial, are higher than the evolutionary rates of BUSCO genes (Fig. 3a), whereas in other squamates the pattern is reversed, with OXPHOS genes appearing more conserved. Longer branches could be associated with the long-branch attraction artifact (LBA) (Felsenstein 1978), but this possibility has already been excluded, at least for mitochondrial OXPHOS subunits (Castoe et al. 2009). This suggests that the high number of parallel substitutions has indeed misled the phylogenetic reconstruction, resulting in mitonuclear discordance (Castoe et al. 2009). They estimated that 113 convergent changes distributed across all 13 mitochondrial protein-coding genes shaped the phylogenetic affinity between snakes and agamid lizards, two lineages separated by more than 160 million years of evolution (Kumar et al. 2017). Similarly, here we found that 11 out of 13 mtOXPHOS showed convergent amino acid substitutions between these two lineages. We might suppose that our methodological approach does not fit well with extremely short sequences like the ones of ATP8 and NADH4L, the only two subunits with different outcomes. COX1 reported the highest number of convergent mutations, coherently with its in-depth analysis (Castoe et al. 2008). Similarly, CYTB, NADH1, 2, 3, and NADH5 reported a higher number of convergent mutations compared to the number of convergent mutations calculated on other pairs of lineages (Fig. 4a).

According to our results, convergent evolution between agamids and snakes is not restricted only to mitochondrial genes. About 30% of nucOXPHOS subunits reported a rate of convergent substitutions significantly higher than the rate of other pairs of branches (Fig. 4b). We suspected that the broad evolutionary returning of the oxidative phosphorylation mechanism in snakes and agamids was beyond the innermost parts of complexes and beyond just containing the catalytic sites, extending to various nucOXPHOS. This substantial percentage has confirmed our predictions. Although the signal was detected in both contact and core nuclear subunits, as well as supernumerary subunits, there is a clear prevalence of convergent substitutions among nucOXPHOS proteins that physically interact with mitochondrial ones (18 out of 25). No convergences were detected in CII, the only complex that does not have mitochondrial subunits. Other subunits play key roles in the OXPHOS system, even if not directly in contact with mitochondrial subunits. For example, among CIV subunits, COX5A constitutes the first initial COX subassembly by association with COX4 (Vidoni et al. 2017). Similarly, COX11 is a copper-binding protein required for Cu incorporation into the Cu_B_ site of cytochrome c oxidase (Banci et al. 2004). NDUFA4, a subunit initially listed among CI proteins, is the last one to be incorporated in the CIV. While it does not seem essential for the correct CIV assembly, NDUFA4 plays a crucial role in the biogenesis of COX, facilitating its activation (Pitceathly et al. 2013). In this context, OXPHOS coevolution would be more driven by a common selective pressure on core subunits than the physical contact between nuclear and mitochondrial subunits. The OXPHOS pathway appears to be particularly susceptible to environmental constraints, with positive selection and changes in the amino acid sequences of mitochondrial subunits potentially accompanying new metabolic requirements (Grossman et al. 2004; Shen et al. 2010; Yang et al. 2014). For example, multiple lineages of livebearing fishes (Poeciliidae) showed an impressive convergence for mtOXPHOS due to the exposition of hydrogen sulfide springs (H_2_S, that causes the inhibition of mitochondrial ATP production). The consequences were visible as repeated modifications of the same physiological pathways, genes, and codons associated with mitochondrial function (Greenway et al. 2020). But there are other cases of adaptive convergent amino acid changes due to shared physiochemical constraints on mtOXPHOS (Escalona et al. 2017; Guo et al. 2018; Elbassiouny et al. 2020). Thus, it is not surprising that the proposal of a metabolic barrier separating snakes from other squamates is driven by a large episodic burst of adaptive change in snake metabolism, feeding habits, and mitochondrial core subunits in Serpentes, as suggested by Castoe and colleagues (2008, 2009).

While the massive support towards convergences between interacting proteins could serve as a hint supporting the hypothesis of nuclear or mitochondrial compensation, this case could also be viewed from another perspective, given its characteristics. If adaptive changes acted on specific sites of mtOXPHOS subunits genes in snakes and agamids (Castoe et al. 2008; Castoe et al. 2009), and if we observe markedly elevated evolutionary rates as well as convergent evolution in both mtOXPHOS and nucOXPHOS, then we may be witnessing not only nuclear-mitochondrial compensation (Havird and Sloan 2016; Hill 2019), but also synergistic coevolution between mitochondrial and nuclear subunits to enhance OXPHOS functionality. This would make sense considering the well-documented increase in OXPHOS functionality in snakes, which would have benefited from mutations that enhance fitness (Castoe et al. 2008). Synergic coevolution posits that beneficial substitutions in one genome enhances successive beneficial changes in the other genome. This scenario is much more coherent with our data in comparison to the hypothesis of solely nuclear compensation to counterbalance the negative effect of the high mitochondrial mutational rate (Sloan et al. 2017). Thus, it could be a quite different manifestation of mitonuclear coevolution compared to the more literal forms of nuclear or mitochondrial compensation.

## 4.0 CONCLUSIONS

This study provides evidence for a shared evolutionary history between agamids and snakes regarding oxidative phosphorylation metabolism. The sister relationship between these two clades, established using mitochondrial OXPHOS genes, is consistent with findings from a subset of nuclear OXPHOS subunits that physically interact with mitochondrial ones. This alignment is explained by a high level of covariance between the two gene sets, as confirmed by ERC analyses. Although these analyses are crucial for validating the hypothesis of coevolution and protein interaction, they also reveal some intrinsic limitations that require further investigation. Our analysis revealed that nearly all mtOXPHOS subunits and a considerable amount of nucOXPHOS subunits exhibit a significant number of convergent substitution events between agamids and snakes. Interestingly, a substantial portion of these convergent events concerns nuclear proteins that physically interact with mitochondrial subunits, but others are also involved. While these observations might suggest a role for nuclear compensation to balance mitochondrial mutations, it is also conceivable that nuclear mutations contribute to enhanced OXPHOS functionality in both clades. Future studies should focus on individual subunits and specific amino acid changes to thoroughly investigate the different aspects and levels of coevolution within this order, which could serve as a model for the research on mitonuclear coevolution.

## 5.0 METHODS

### 5.1 Mitochondrial OXPHOS annotation

Among the 59 species, complete mitochondrial genomes were available for 36 species on NCBI (Table S1). For the remaining species, we employed various strategies to retrieve and annotate the 13 mitochondrial genes, depending on the available data for each species (Table S1). Initially, we utilized blastn (Blast v2.16.0+; Camacho et al. 2009) to locally align the contigs or scaffolds of each genome assembly against the set of 36 complete mitochondrial genomes to determine whether the mitochondrial genome was already included in the genome assemblies. In 15 species, the mitochondrial genome was one of the contigs included in their genome assembly. These contigs were annotated using MITOS (Bernt et al. 2013). For the remaining species, we searched for whole genome sequencing data produced during the respective genome project. In cases where long PacBio reads were available, we employed MitoHiFi (Allio et al. 2020; Uliano-Silva et al. 2023) to retrieve and annotate the mitochondrial genome. Conversely, when only short Illumina reads were available, we used SPAdes (Bankevich et al. 2012) to assemble the short reads into contigs. To identify mitochondrial contigs, we locally aligned them with blastn against the set of 36 mitochondrial genomes. Subsequently, mitochondrial contigs were annotated using MITOS.

### 5.2 Nuclear OXPHOS annotation

Gene list and nomenclature for nuclear OXPHOS (nucOXPHOS) were sourced from the Kyoto Encyclopedia of Genes and Genomes (KEGG, map00190; Kanehisa and Goto 2000). 18 out of 59 species had accessible proteomes from RefSeq: we retrieved orthogroups associated with nucOXPHOS from OrthoDB v11 (Kuznetsov et al. 2023) and extracted sequences for these species. For the remaining 41 species, Miniprot (Li 2023) was used to align protein sequences from the closest RefSeq species to their assemblies. Given that Miniprot aligns each protein independently, it may introduce errors when slightly divergent paralogs are present: those resulting from redundant annotation of Miniprot were manually inspected and removed. We used Augustus (Stanke et al. 2004) to finally predict the nucOPXHOS gene, selecting the region where Miniprot aligned the protein and giving the Miniprot alignment as hint. Since paralogs were present for at least one species in nearly all nucOXPHOS genes, we inferred the ML tree for each gene using IQ-TREE v2.0.3 (Minh et al. 2020), and we manually inspected every gene tree to select orthologous groups and discern them from paralogous. From the nucOXPHOS dataset we selected a subset of genes, the “contact nucOXPHOS” dataset, which are in physical association with mitochondrially encoded subunits (Fig. 2a). To select this subset of genes we interrogated the literature about OXPHOS complexes (CIV: Richter and Ludwig 2003, CV: Jonckheere et al. 2012; CI: Zhu et al. 2016; CIII: Amporndanai et al. 2018). Additionally, a control was performed using the crystal structures from GeneCards (Stelzer et al. 2016).

The completeness matrices were constructed using RStudio v4.3.1 (R Core Team 2021) with ggplot2 (Wickham 2016) (Fig. S2).

### 5.3 Orthologs annotation

We created a set of orthologous genes to be used in comparative analysis along with mtOXPHOS and nucOXPHOS datasets. The BUSCO v.5.5.0 (Simão et al. 2015) pipeline was run with the Metazoa_odb10 dataset to assess the completeness of our assemblies (Table S1). Species with low BUSCO completeness scores (< 50%) were removed from the dataset. We included all single copy BUSCO genes that were inferred in at least 80% of our species. We made sure that all OXPHOS genes were filtered out from the BUSCO gene dataset.

### 5.4 Phylogenetic analysis and Site-specific Likelihood Support

The amino acid sequences of mtOXPHOS and nucOXPHOS genes were aligned using MAFFT v7.52 (-auto; Katoh and Standley 2013). Alignments were masked using ClipKit (Steenwyk et al. 2020). Nucleotide sequences were aligned and masked according to the amino acid alignments. Orthogroups were concatenated into two matrices corresponding to the two datasets using AMAS.py (Borowiec 2016).

For BUSCO genes, multiple sequence alignment, trimming, and creation of a supermatrix were performed using the “BUSCO_phylogenomics.py” pipeline (McGowan 2023), both for nucleotide and amino acid sequences. For each dataset, we inferred maximum-likelihood (ML) trees from the partitioned nucleotide and amino acid datasets using IQ-TREE (Minh et al. 2020) with 1,000 ultrafast bootstrap replicates (Hoang et al. 2018). We used Modelfinder to perform model selection (Kalyaanamoorthy et al. 2017) and we specified the MFP+MERGE option (Chernomor et al. 2016). ML analysis was also conducted on the contact nucOXPHOS subset. Bayesian inference was performed using PhyloBayes (Lartillot et al. 2009). The phylogenetic trees were plotted using iToL (Interactive Tree of Life) (Letunic and Bork 2024). Site-specific likelihood support (SSLS) analysis was conducted on the nucOXPHOS alignment to assess the site-specific support for alternative topologies (i.e., “mitochondrial” and “nuclear” topologies). SSLSs associated to mtOXPHOS and nucOXPHOS topologies were retrieved performing RAxML with the “-f G” option under the GTRGAMMAI model (Stamatakis 2014).

### 5.5 Branch lengths comparisons and Evolutionary Rate Correlations

The branch lengths of our four datasets (mtOXPHOS, nucOXPHOS, contact nucOXPHOS and BUSCO genes) were optimized on a consensus tree (Burbrink et al. 2020) using the “-f e” option of RAxML (Stamatakis 2014). Root-to-tip distances were retrieved using the distRoot function (tips = “all”, method = “patristic”; R package: adephylo; Jombart et al. 2010)). Subsequently, all lengths were scaled by dividing each value by the sum of all root-to-tip distances, to compared the evolutionary rates of mtOXPHOS, nucOXPHOS and BUSCO genes across the 56 OTU. Evolutionary rate correlations (ERC) were used to investigate the coevolution between datasets. Since correlations between branch lengths are exposed to phylogenetic bias (Phylogenetic Independent Contrast; Felsenstein 1985), we developed a method to avoid it and test correlation between branch lengths, based on the distribution of Spearman’s ρ values. We generated four random and independent subsets of the BUSCO genes, consisting of two sets of 73 genes (i.e., number of nucOXPHOS genes) and two sets of 13 genes (i.e., number of mtOXPHOS genes). We repeated the extraction 1000 times. Half of the resulting subsets were used for the normalization, while the remainder was employed for correlation testing (see Fig. S6). After normalization, the cor.test R function was performed between branch lengths of i) mtOXPHOS and nucOXPHOS, ii) nucOXPHOS and BUSCO genes, iii) mtOXPHOS and BUSCO genes, and iv) mtOXPHOS and contact subunits of nucOXPHOS. Density curves of Spearman’s ρ values are plotted in Figure 3b (R packages - ggplot2: Wickham 2016, hrbrthemes: Rudis et al. 2019, dplyr: Wickham et al. 2023, tidyr: Wickham et al. 2024, viridis: Ganier et al. 2024).

### 5.6 Codon convergences

We used MEME (Murrell et al. 2012) to infer all the substitutions along the Squamata species tree, whose topology was built following the relationships of the Squamata tree (Burbrink et al. 2020). Using custom-made python scripts, we retrieved all the substitutions occurred in all branches. Then, we compared all possible independent branches (i.e., when the group of OTUs subtended by one branch does not intersect with the group of OTUs subtended by the other branch), excluding terminal ones, counting the number of convergent and divergent substitutions. We consider a substitution when it occurred on both branches in the same codon, and we defined it convergent when the mutation led to the same amino acid, while we defined it divergent when it led to different amino acids. Although we analyzed all possible pairs of independent branches, we specifically highlighted the combinations between basal groups, focusing on pairings between Agamidae and the main outgroup taxa, as controls. The same approach was then applied to combinations within Serpentes. A schematic overview of the workflow was presented in Fig. 3c.

## Data Availability

Custom tailored Python, bash and R scripts written for all the analyses have been uploaded on a public GitHub repository (https://github.com/oscarwallnoefer/Squamata) and are available by O.W. upon reasonable request.

## Author contributions

A.F. and F.P. conceived the project; A.F. and O.W. analysed the data and wrote the first draft of the article; O.W. designed and prepared the figures; F.P. and M.P. supervised the project; M.P secured funding for the project. All authors accepted and contributed to the final version of the article.

