## Supplementary material for "Evidence of convergent evolution in the nuclear and mitochondrial OXPHOS subunits across the deep lineages of Squamata": S3_TreePlots.pdf

**SUPPLEMENTARY MATERIALS**

**Figure S3.** Phylogenetic trees inferred from the BUSCO genes, mitochondrial OXPHOS and nuclear OXPHOS datasets. **a, b)** BUSCO trees obtained through Maximum Likelihood analysis (IQ-TREE) on nucleotide and amino acid sequences. **c, d, e, f)** mtOXPHOS trees obtained through Maximum Likelihood analysis (IQ-TREE) and Bayesian analysis (PhyloBayes) on nucleotide and amino acid sequences. **g, h)** nucOXPHOS trees obtained through Maximum Likelihood analysis (IQ-TREE) on nucleotide and amino acid sequences. **i)** Contact nucOXPHOS tree obtained through Maximum Likelihood analysis (IQ-TREE) on nucleotide sequences.
Node supports are shown when <99 (ML) or <0.99 (Bayesian).

**a)**
**BUSCO genes**
**ML, nt**

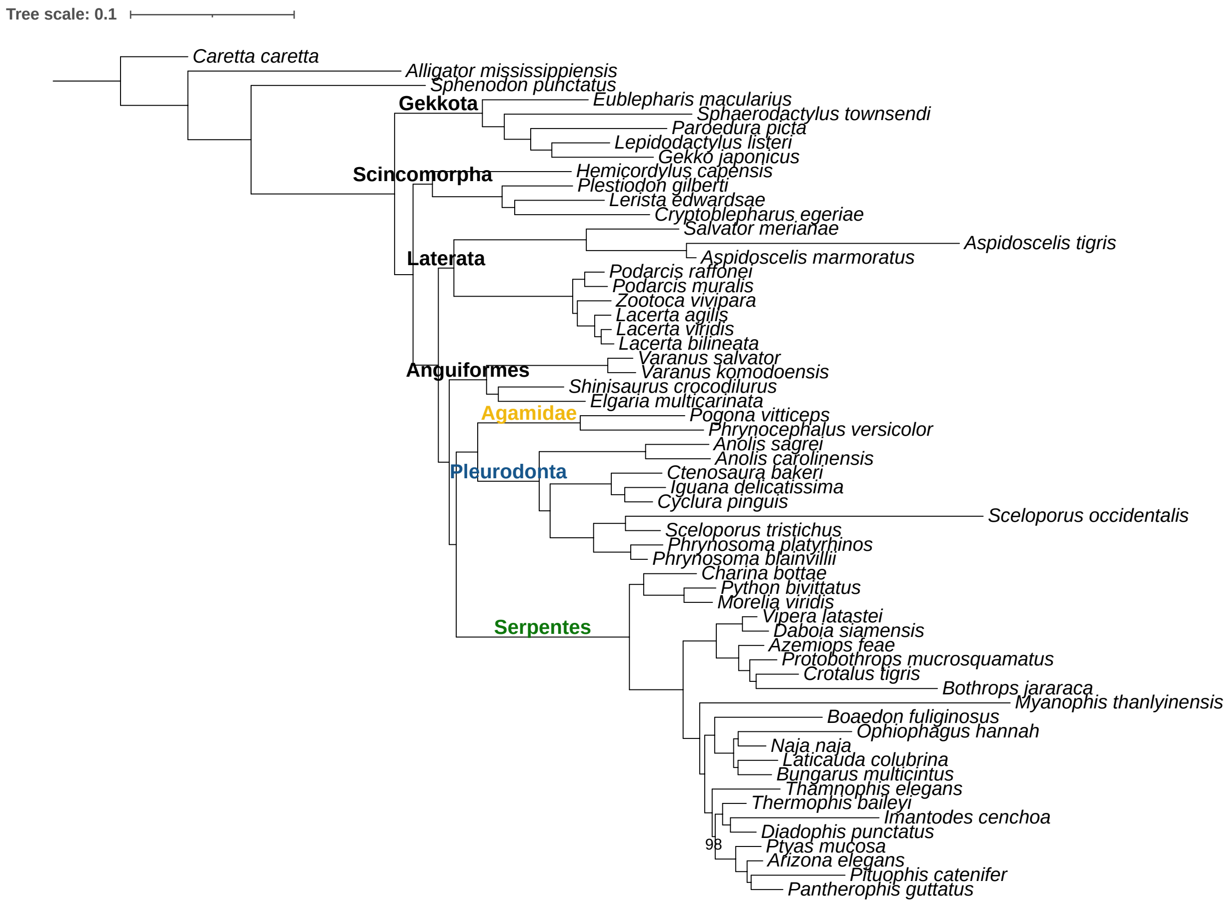

**b)**
**BUSCO treefile**
**ML, aa**

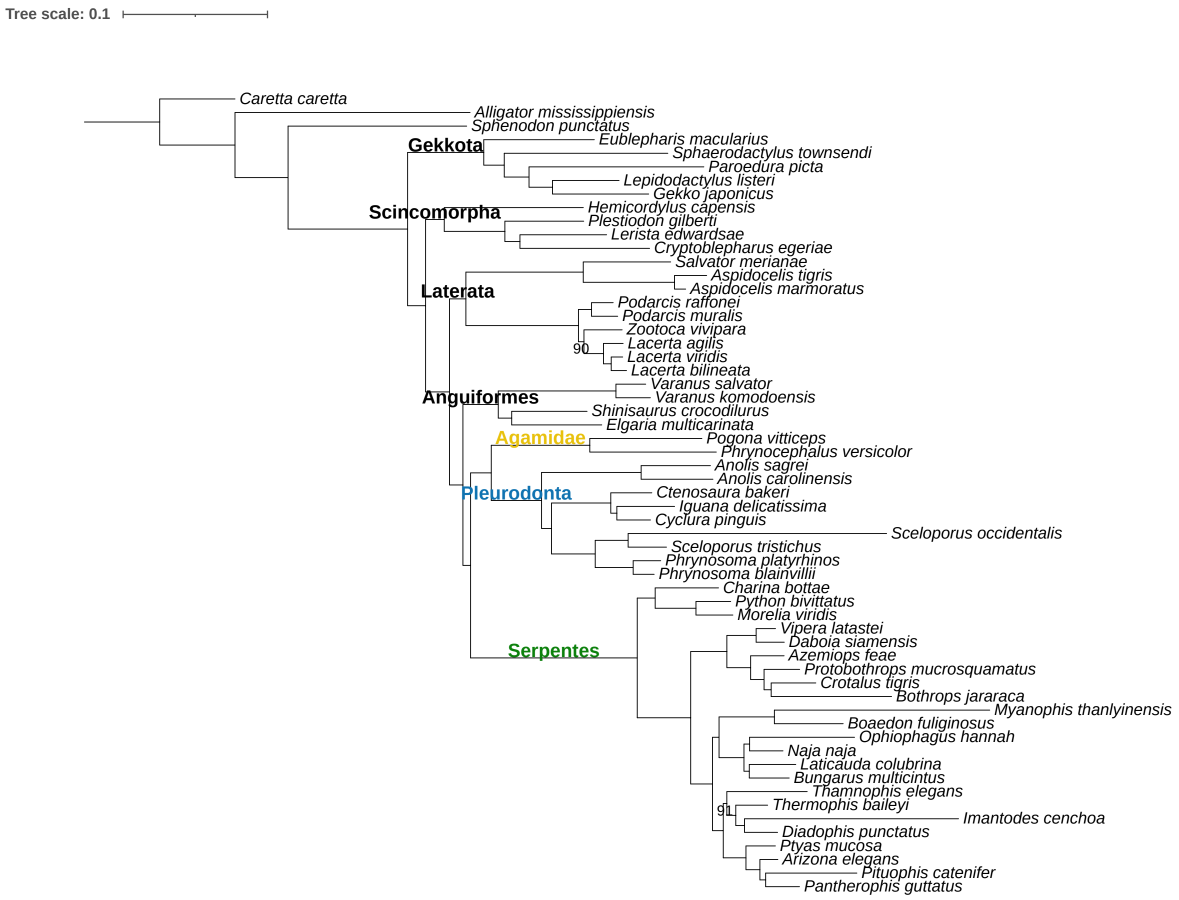

**c)**
**mtOXPHOS genes**
**ML, nt**

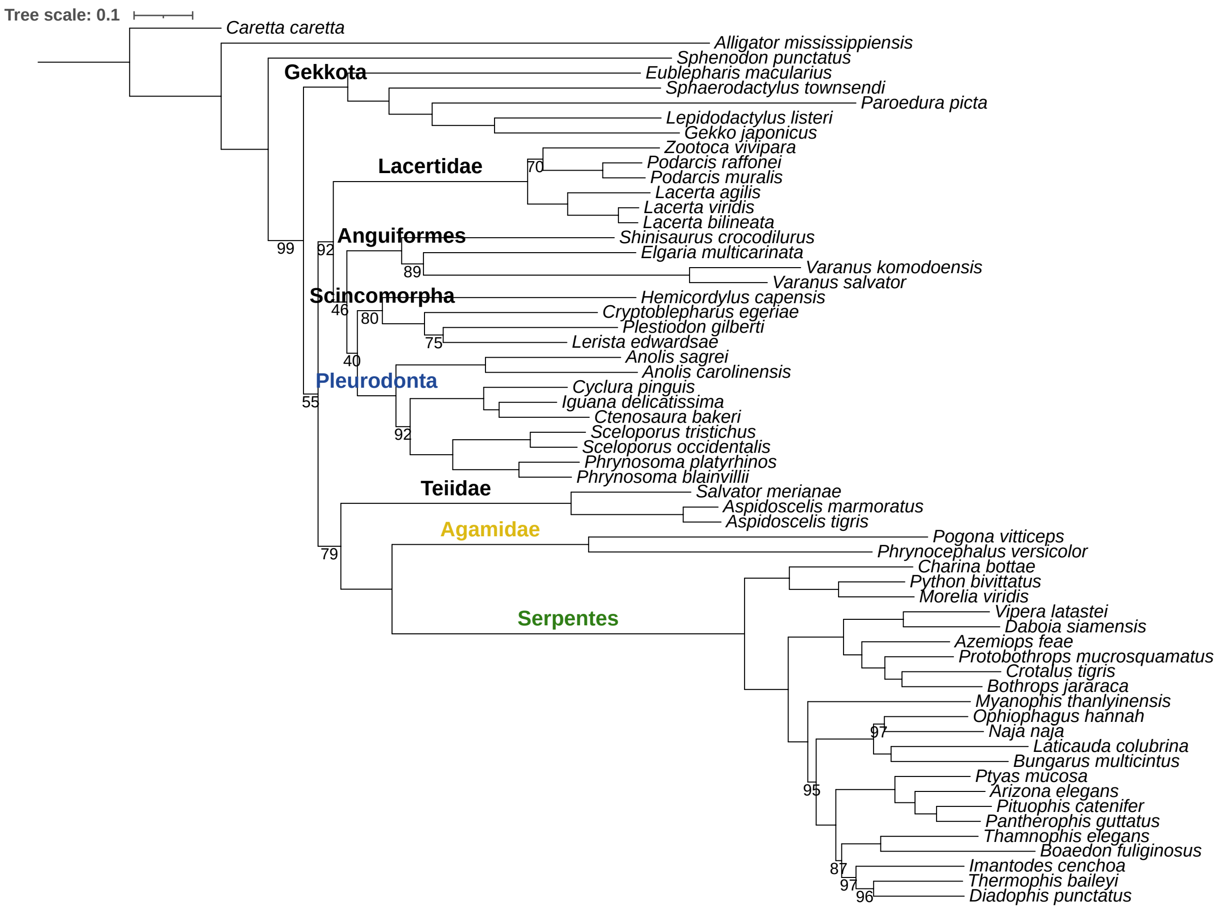

**d)**
**mtOXPHOS genes**
**Phylobayes, nt**

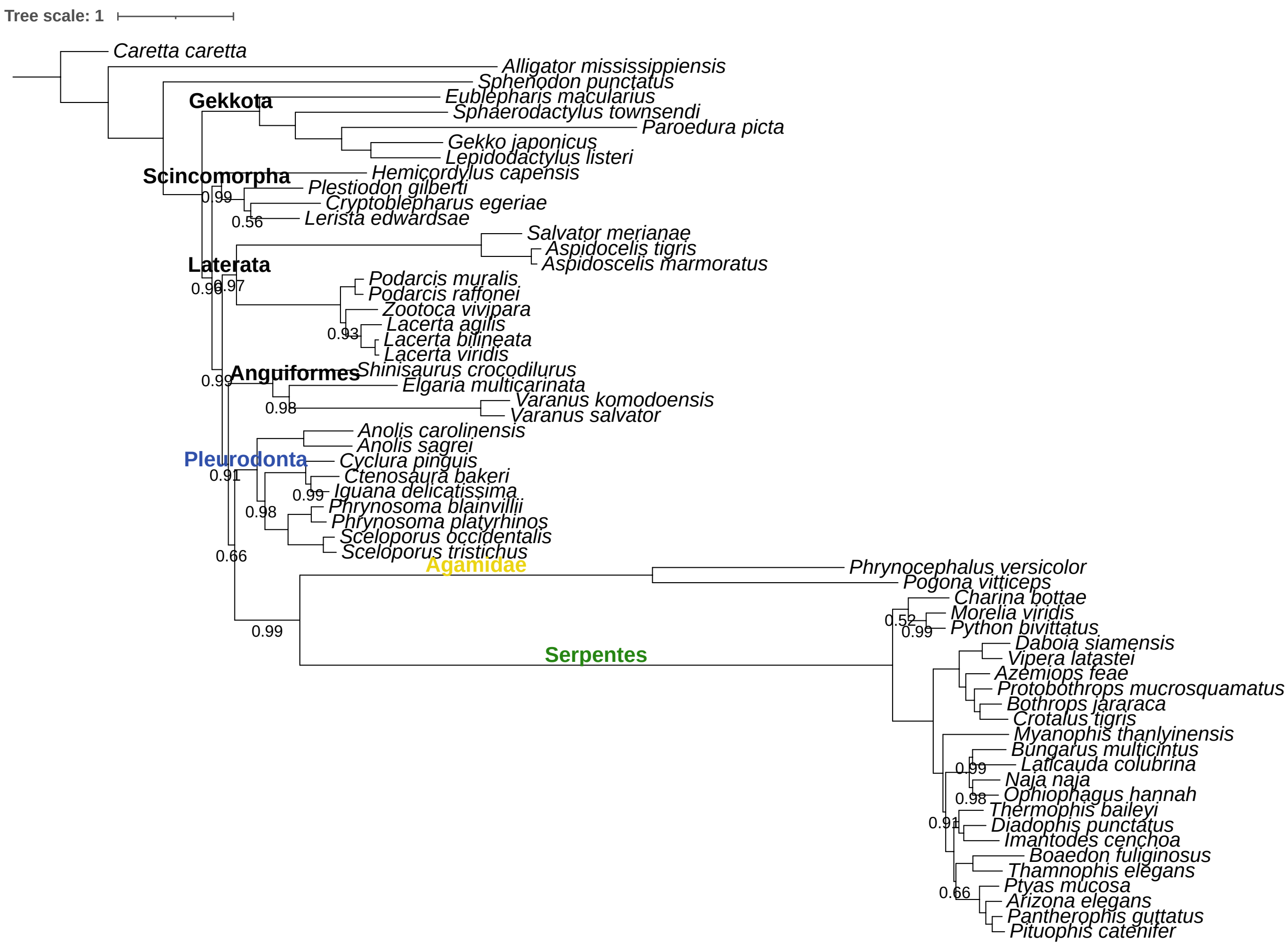

**e)**
**mtOXPHOS genes**
**ML, aa**

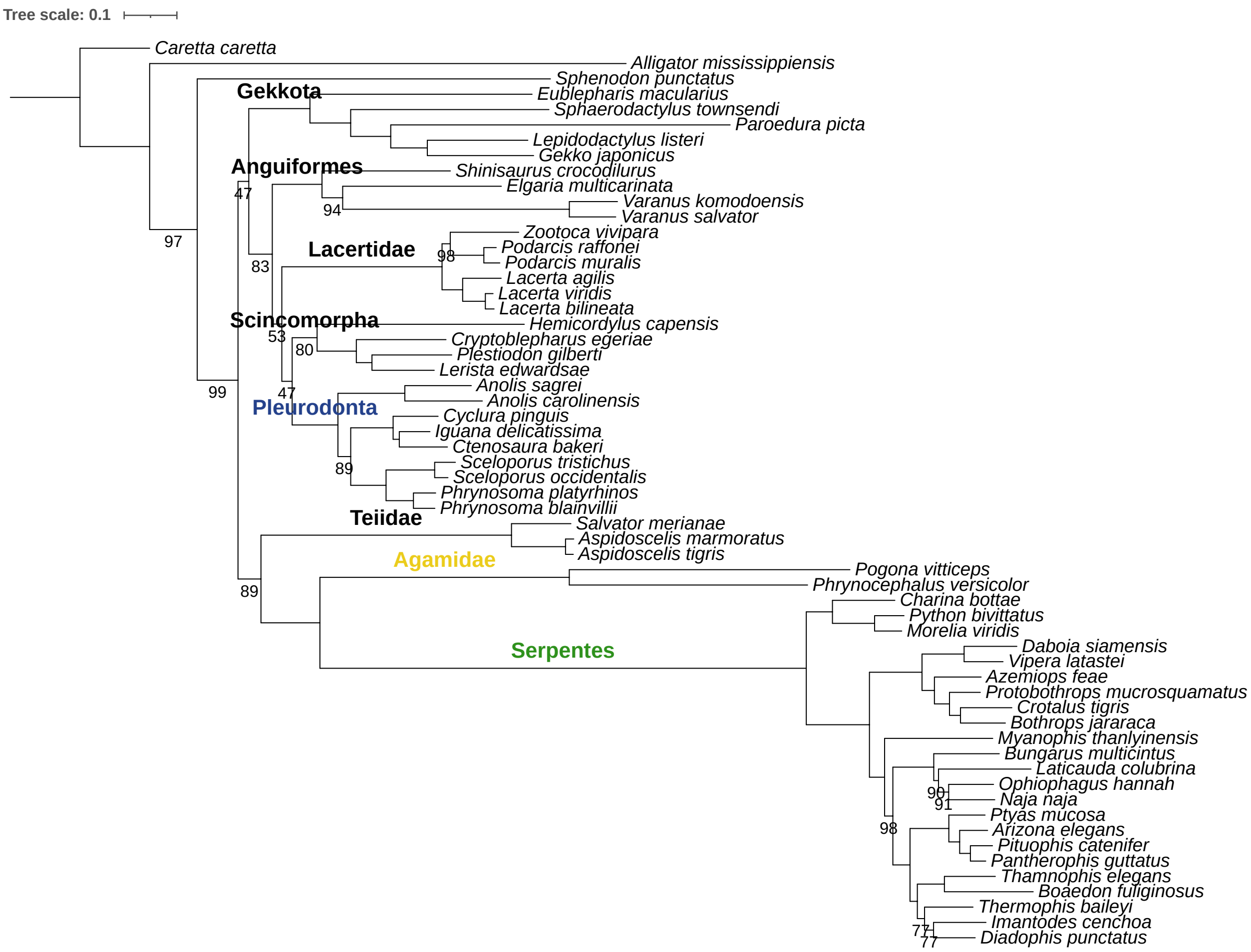

**f)**
**mtOXPHOS genes**
**Phylobayes, aa**

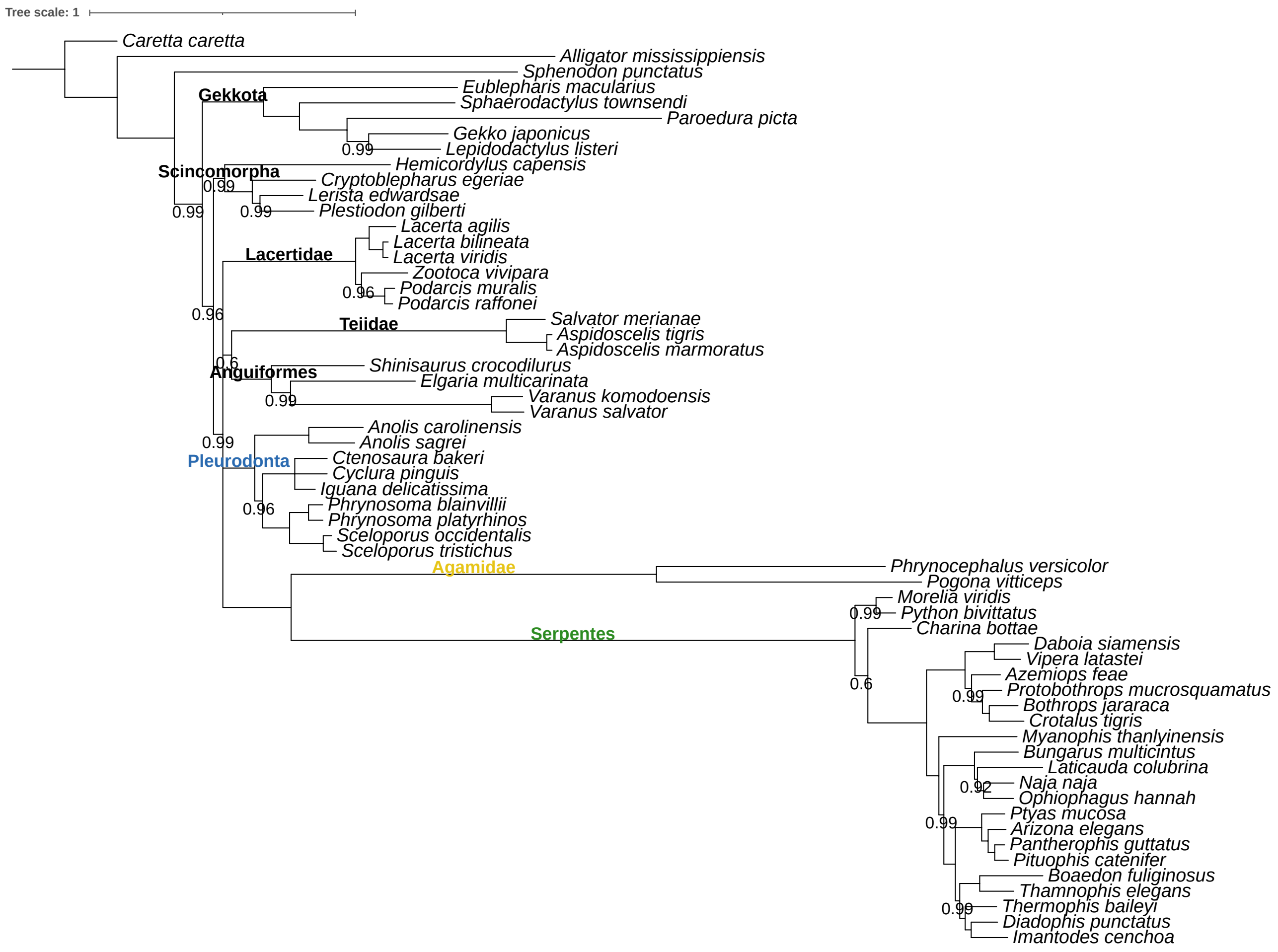

**g)**
**nucOXPHOS genes**
**ML, nt**

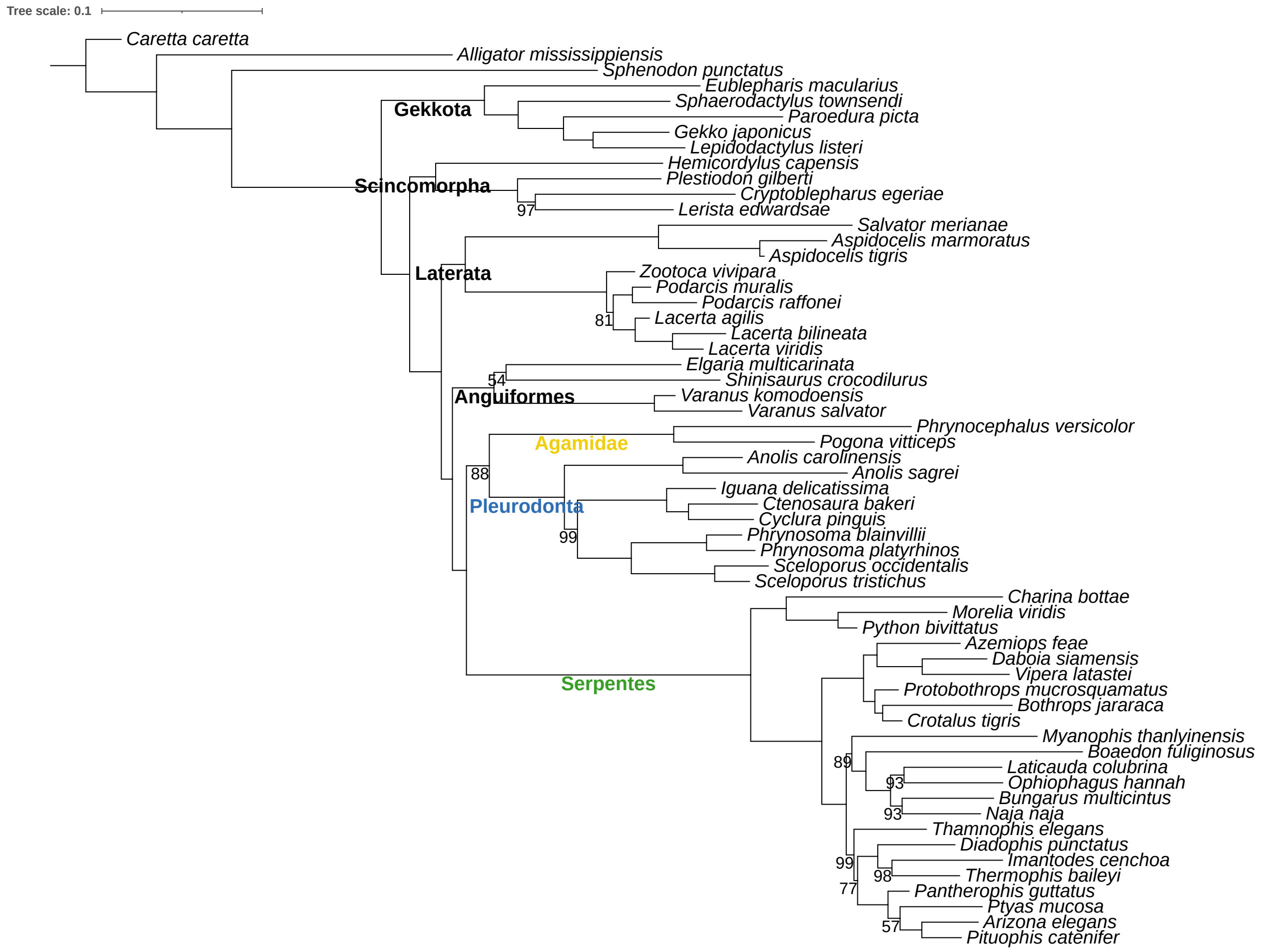

**h)**
**nucOXPHOS genes**
**ML, aa**

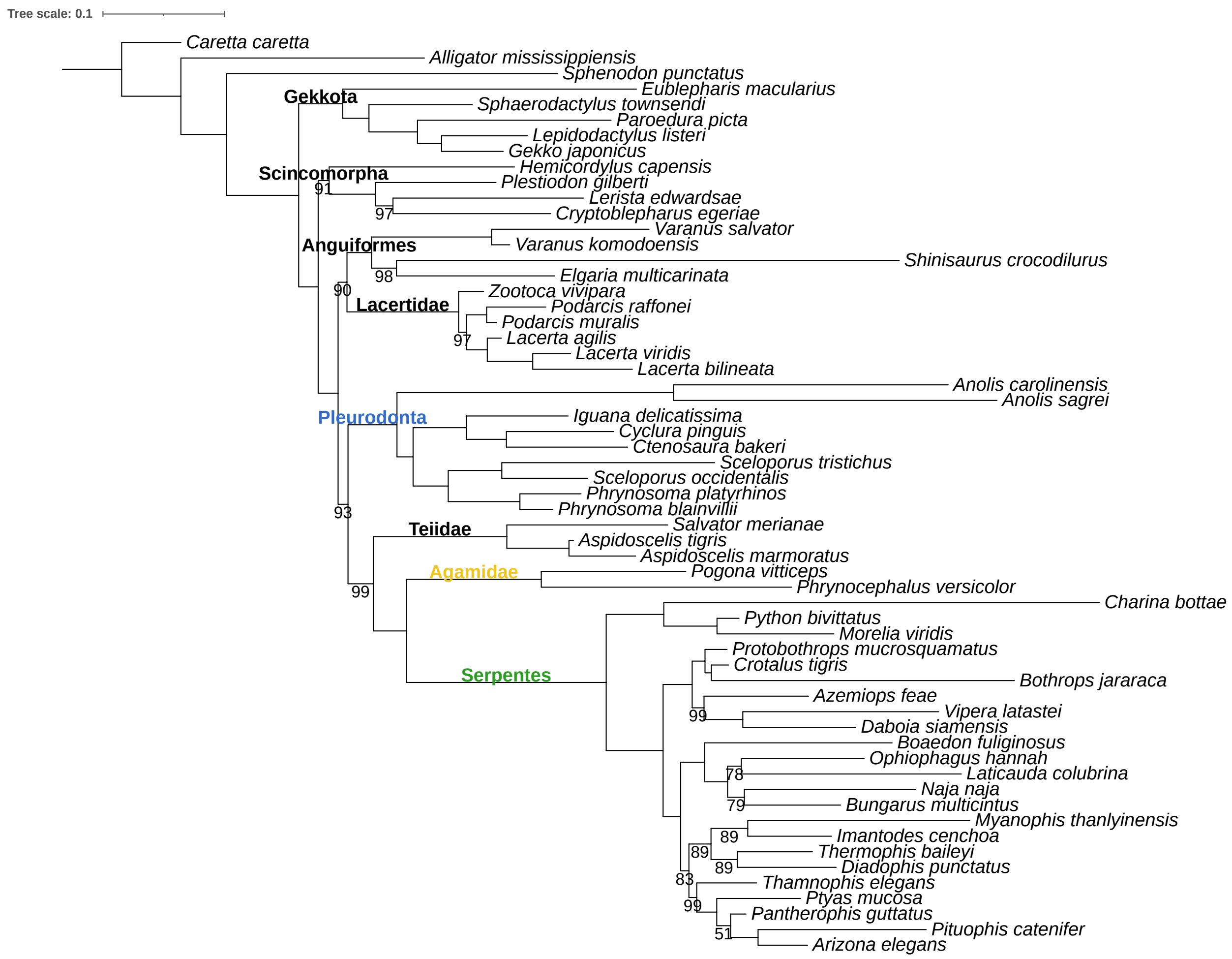

i)
contact nucOXPHOS genes
ML, nt

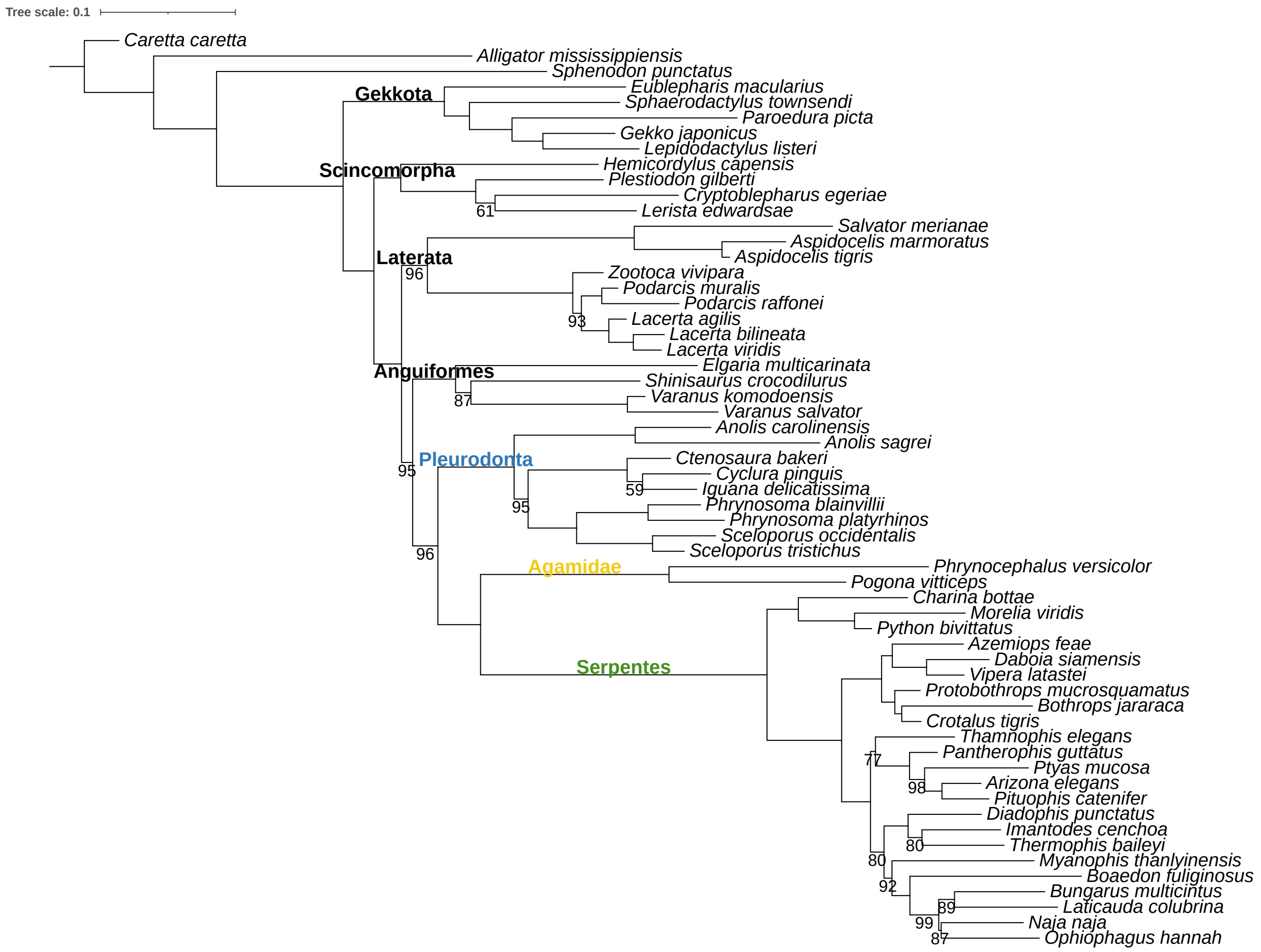
