## Supplementary material for "Evidence of convergent evolution in the nuclear and mitochondrial OXPHOS subunits across the deep lineages of Squamata": S4_Saturation_Test.pdf

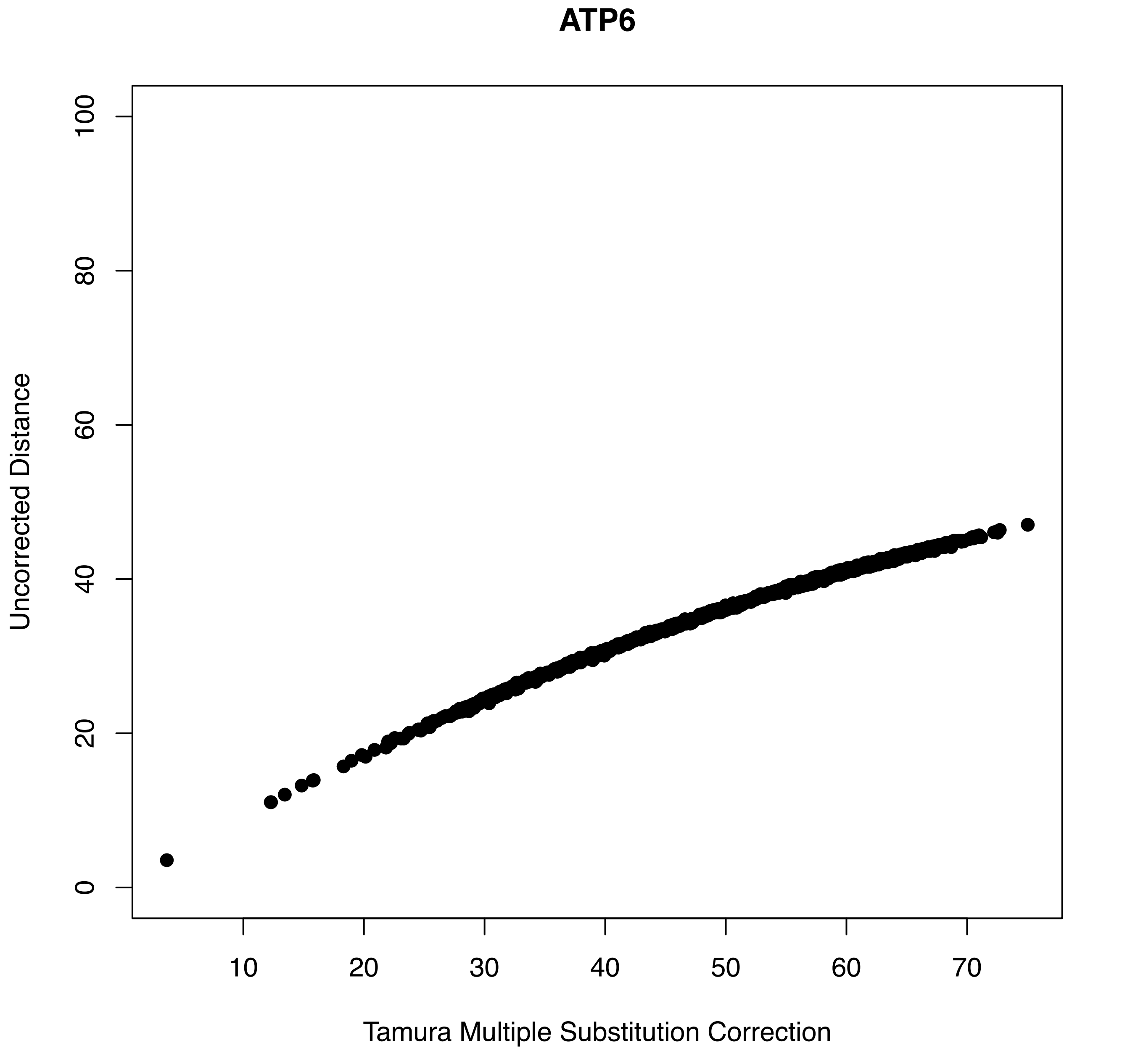
ATP6
Uncorrected Distance
0
20
40
60
80
100
10
20
30
40
50
60
70
Tamura Multiple Substitution Correction

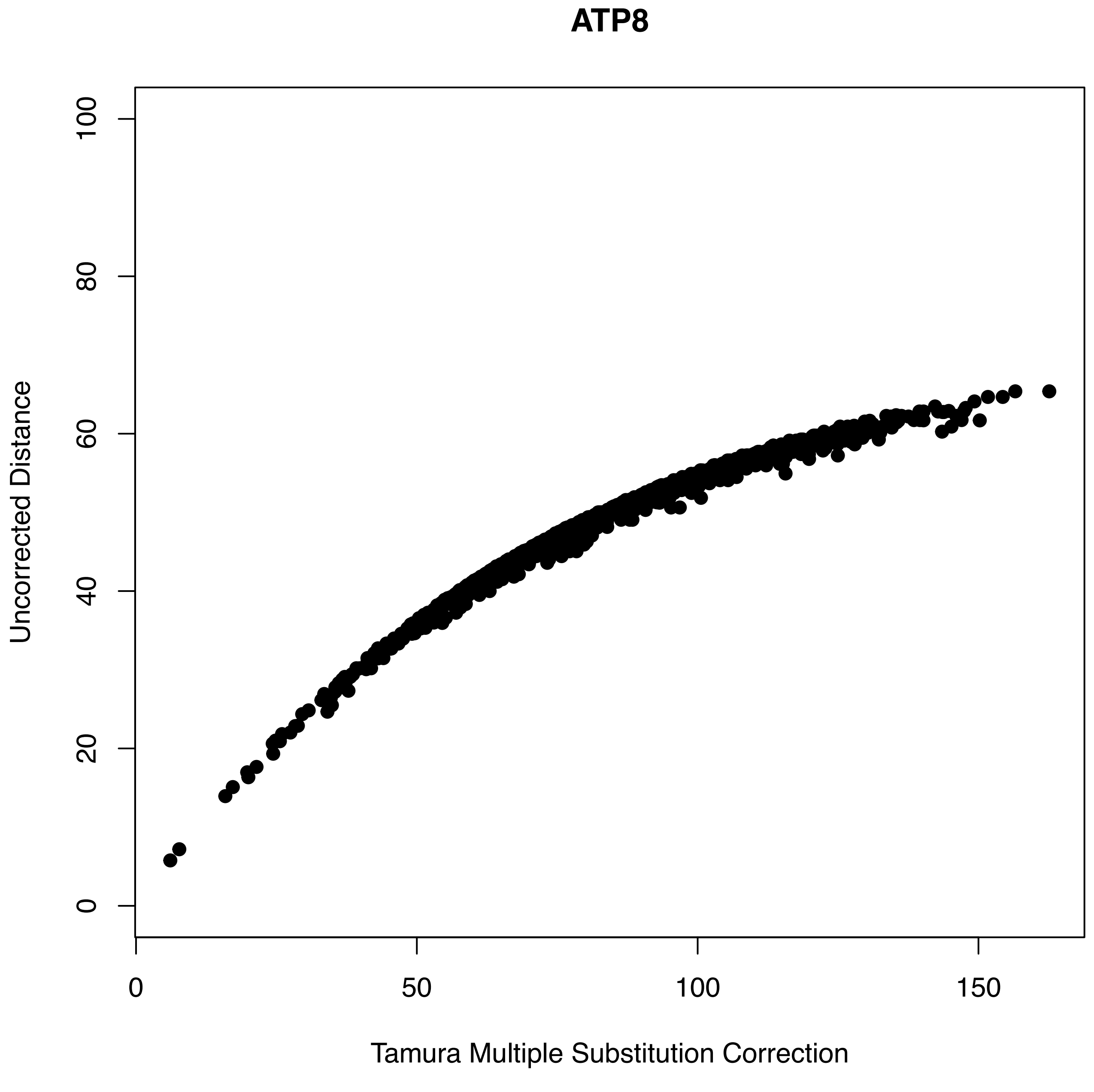
ATP8
Uncorrected Distance
0
20
40
60
80
100
0
50
100
150
Tamura Multiple Substitution Correction

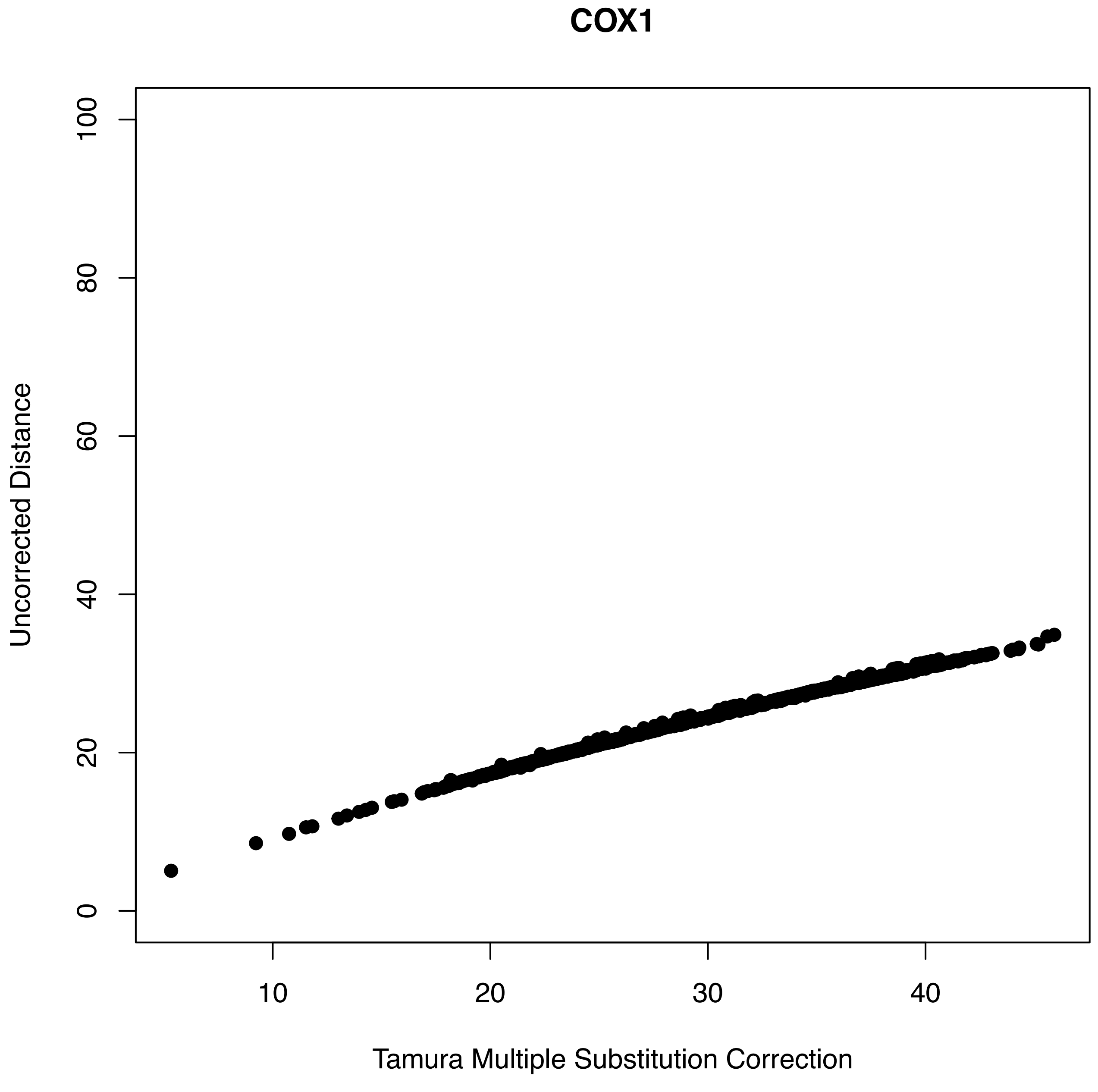
COX1
Uncorrected Distance
0
20
40
60
80
100
10
20
30
40
Tamura Multiple Substitution Correction

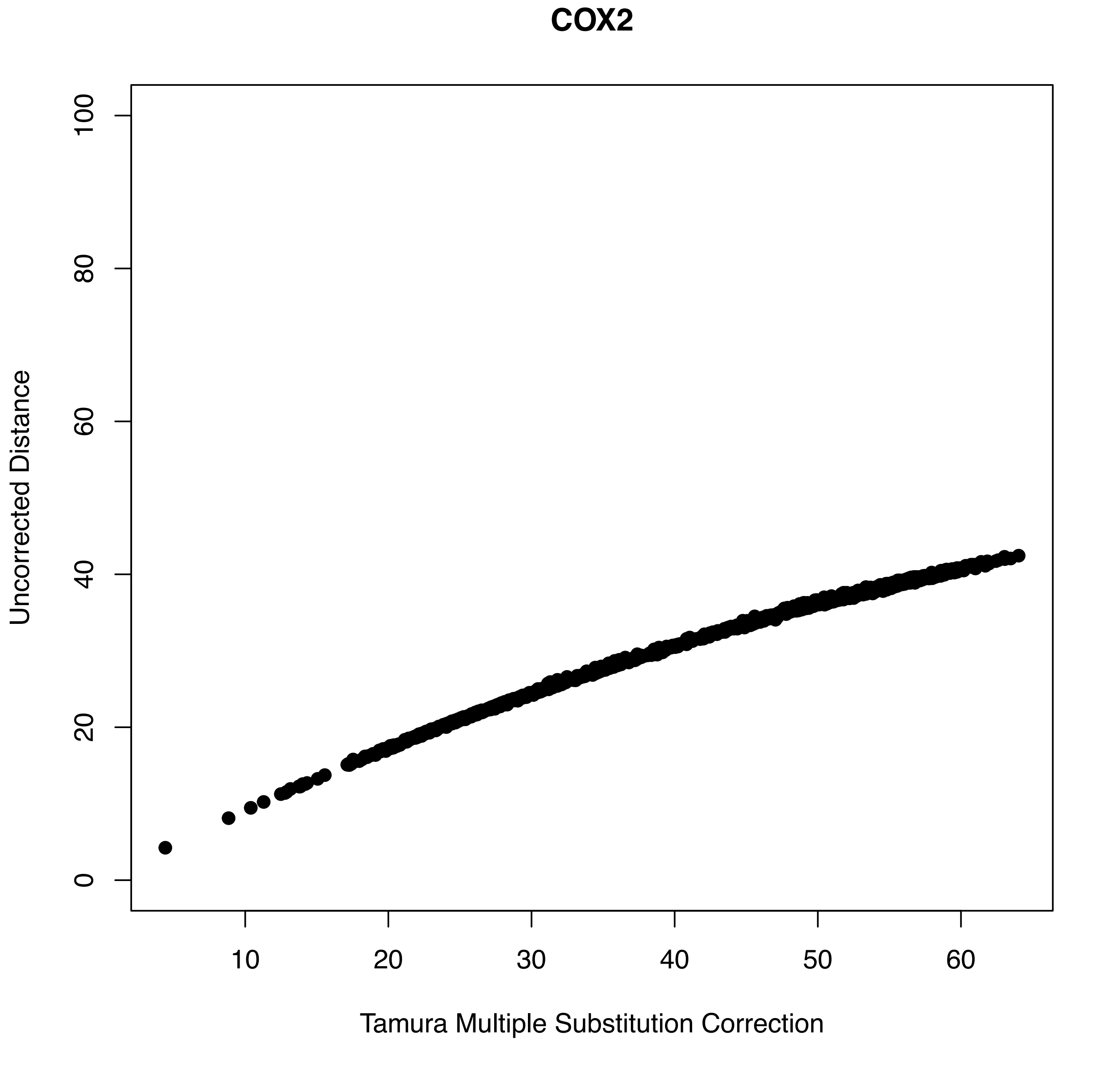
COX2
Uncorrected Distance
0
20
40
60
80
100
10
20
30
40
50
60
Tamura Multiple Substitution Correction

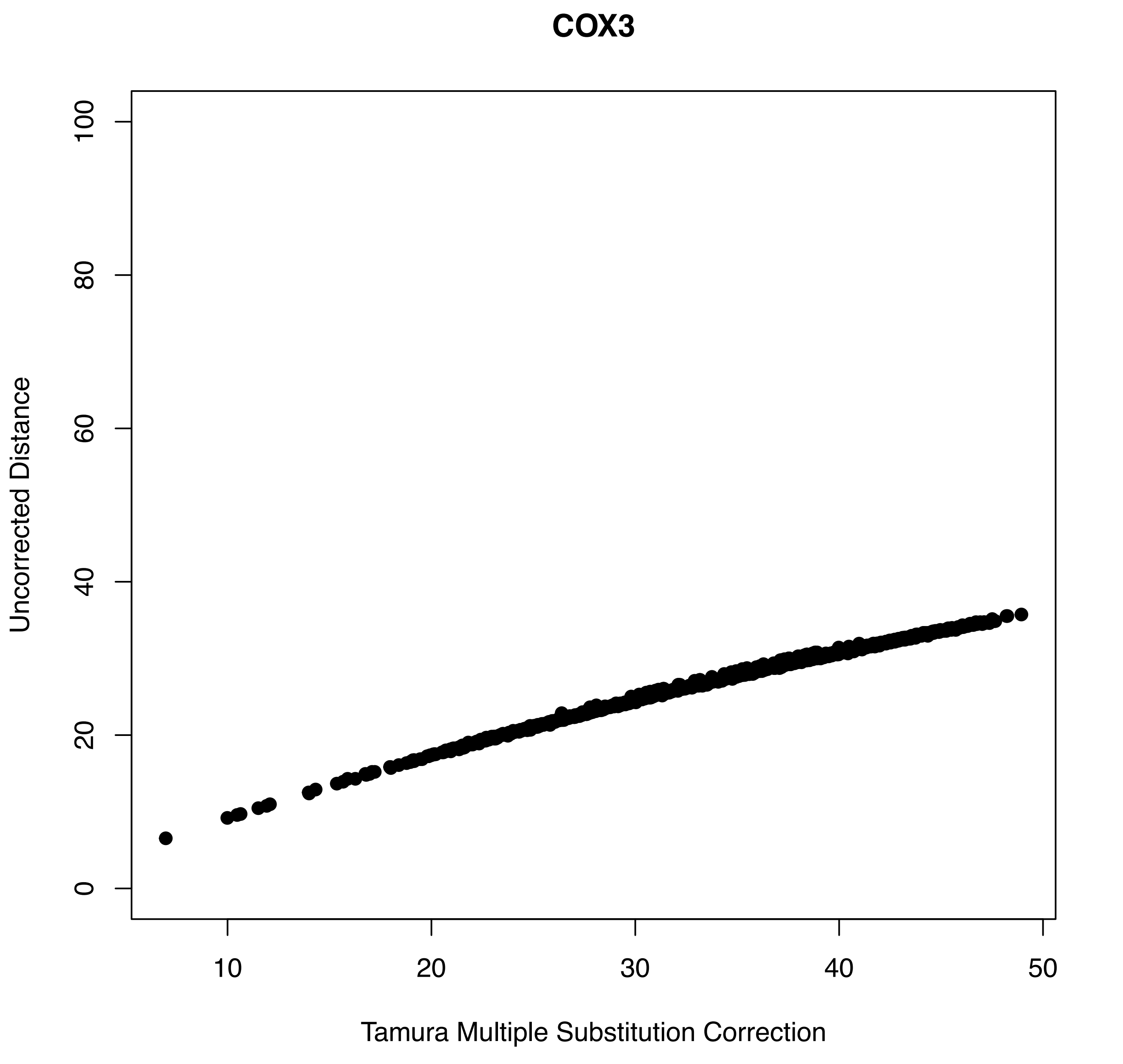
COX3
Uncorrected Distance
0
20
40
60
80
100
10
20
30
40
50
Tamura Multiple Substitution Correction

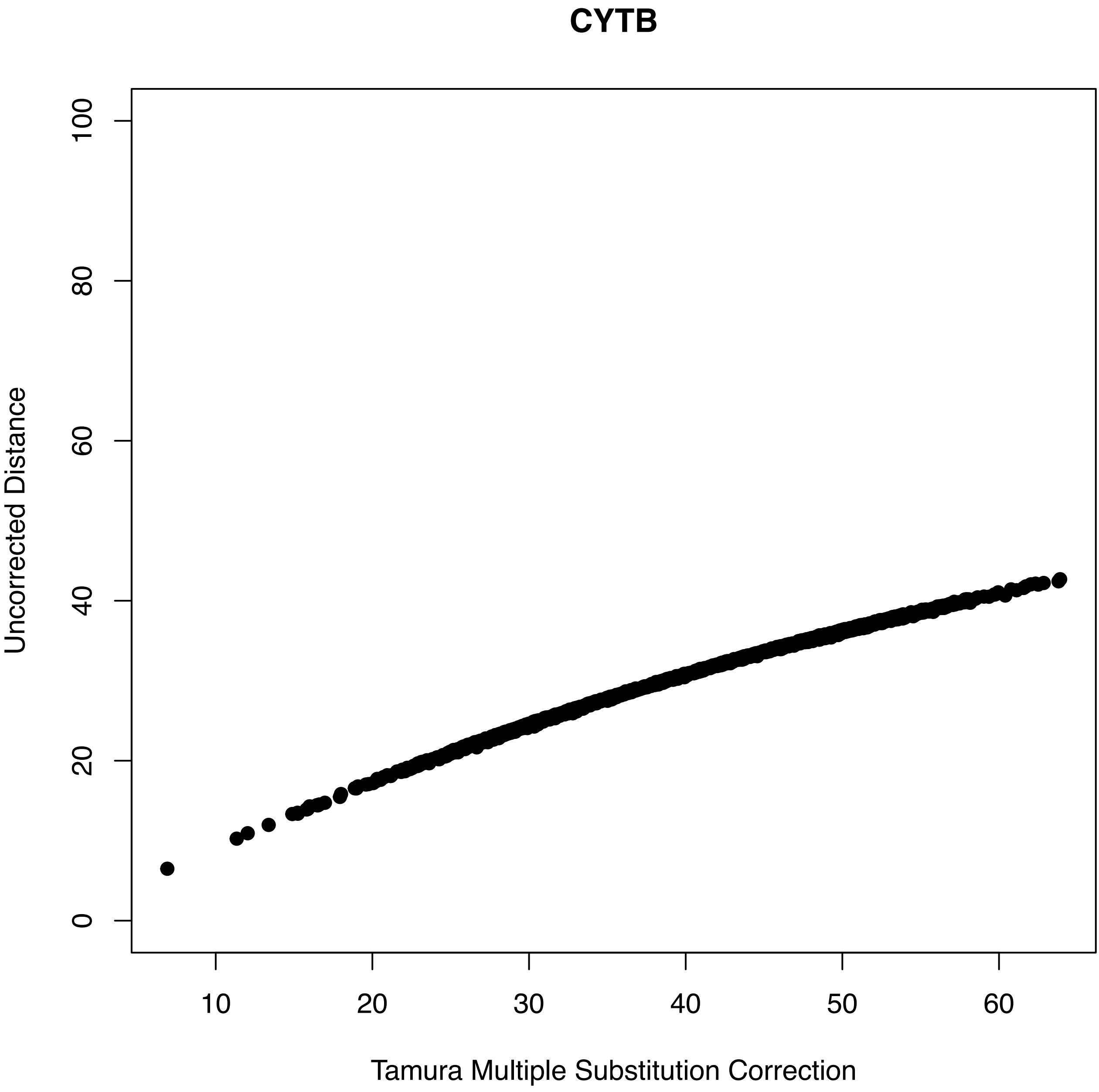
CYTB
Uncorrected Distance
0
20
40
60
80
100
10
20
30
40
50
60
Tamura Multiple Substitution Correction

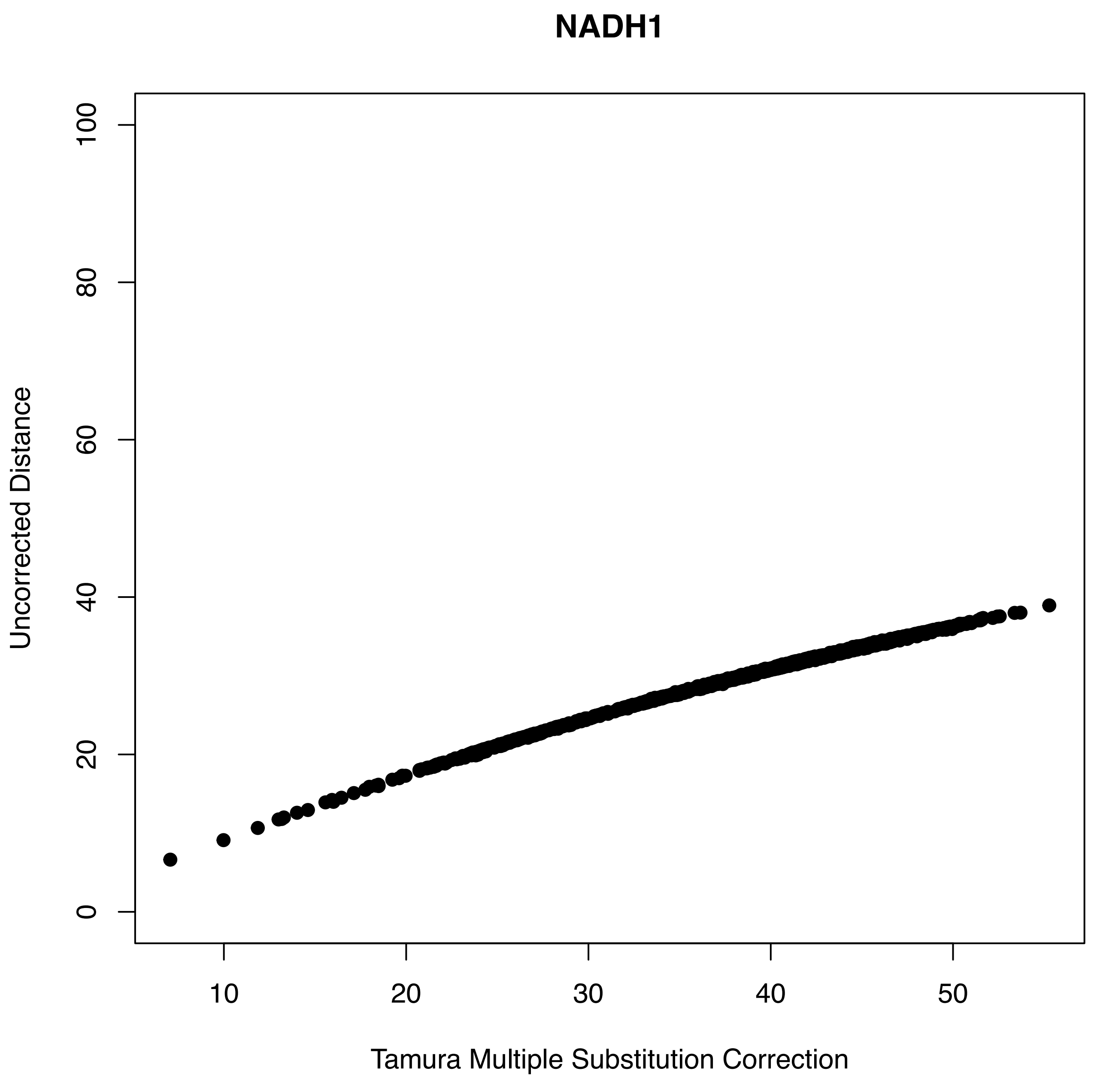
NADH1
Uncorrected Distance
Tamura Multiple Substitution Correction
0
20
40
60
80
100
10
20
30
40
50

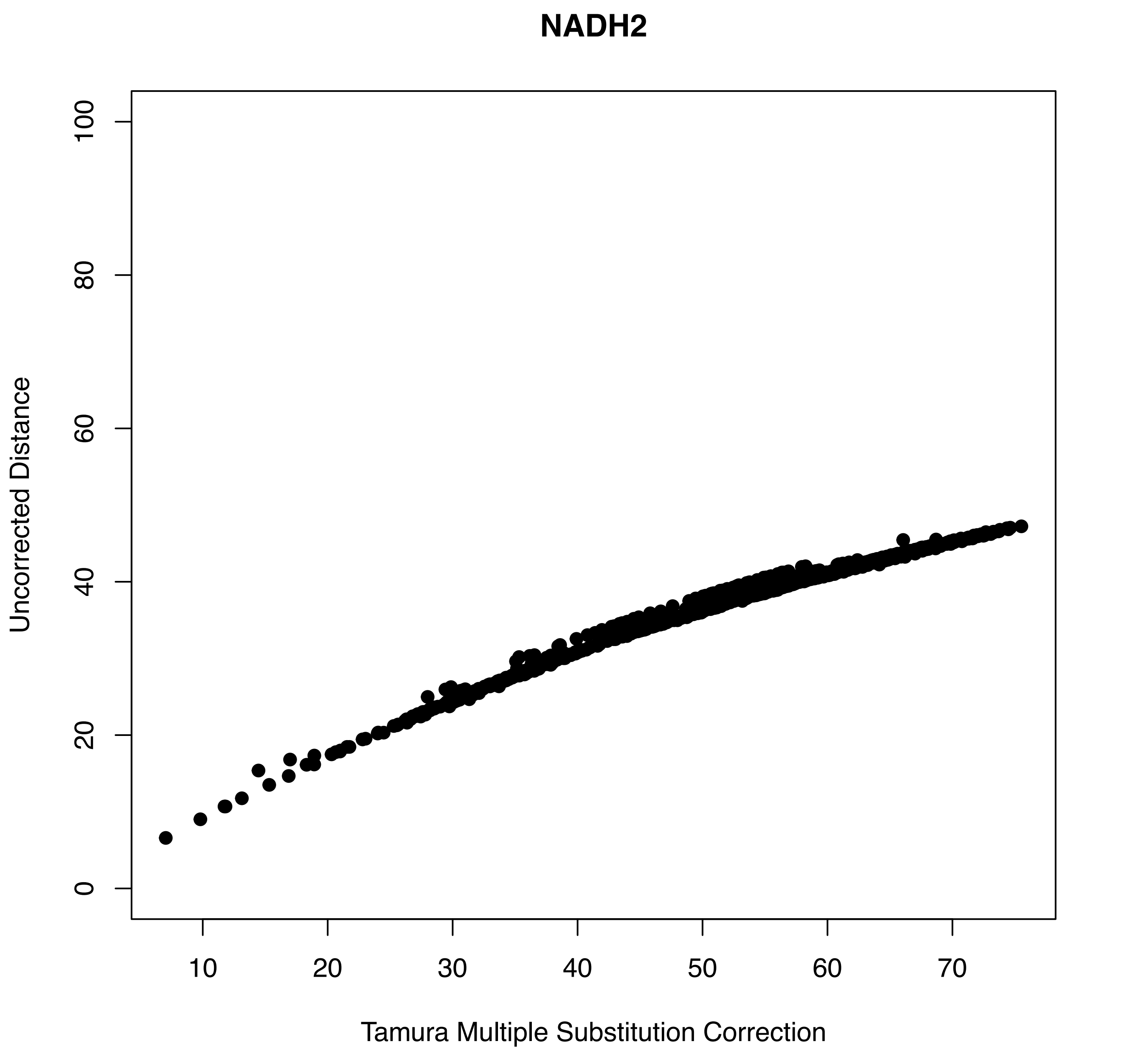
NADH2
Uncorrected Distance
0
20
40
60
80
100
10
20
30
40
50
60
70
Tamura Multiple Substitution Correction

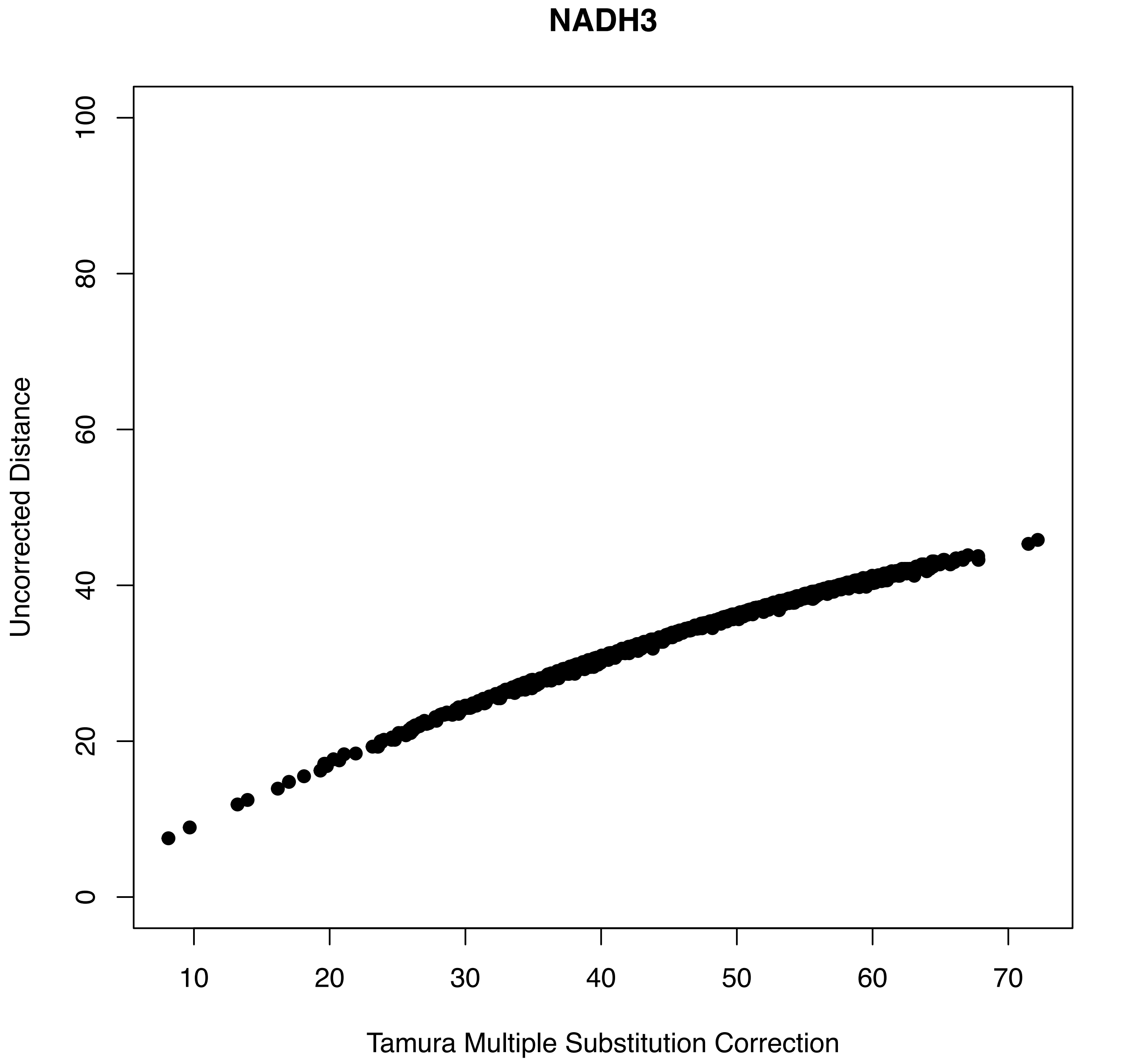
NADH3
Uncorrected Distance
0
20
40
60
80
100
10
20
30
40
50
60
70
Tamura Multiple Substitution Correction

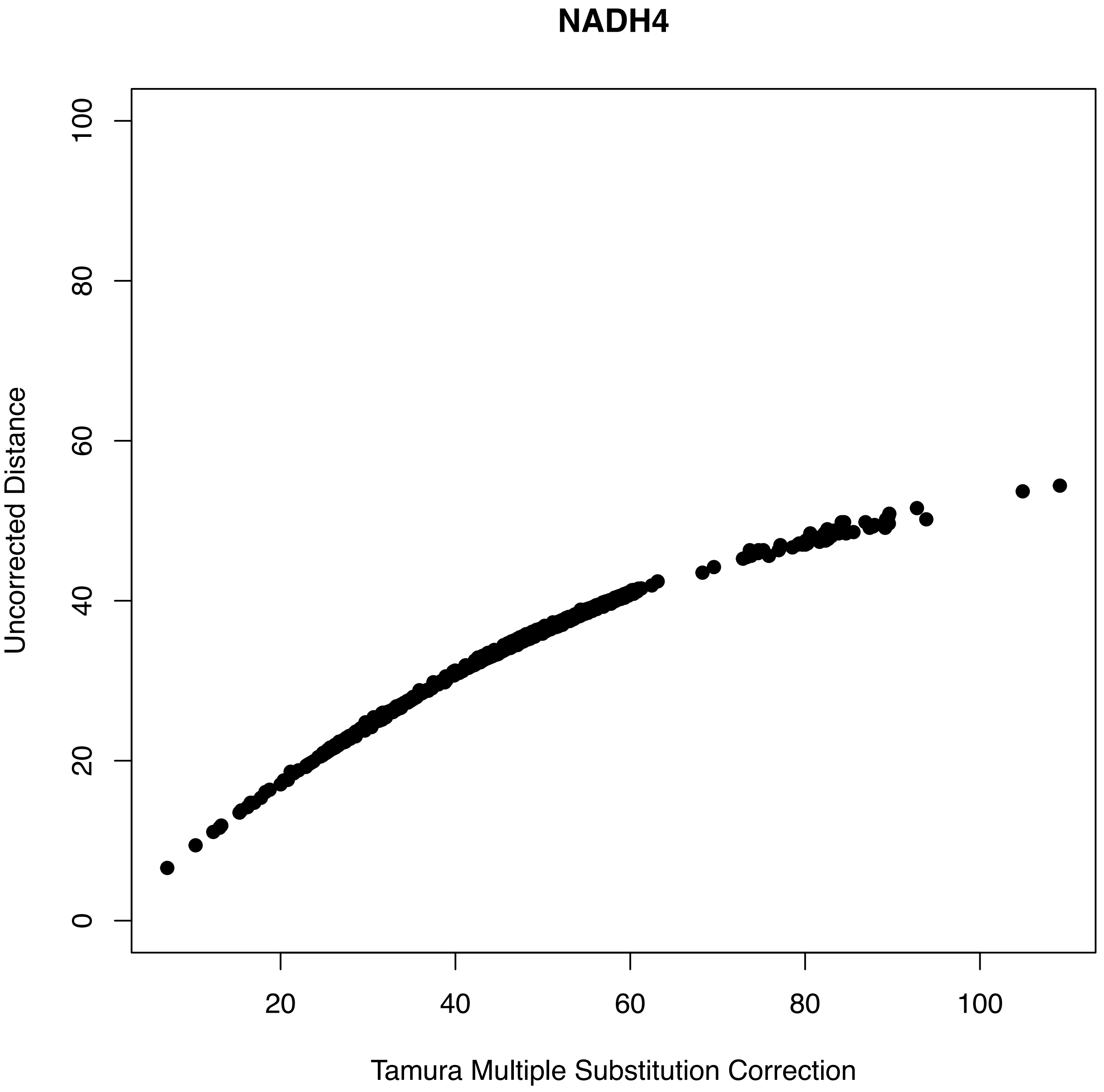
NADH4
Uncorrected Distance
Tamura Multiple Substitution Correction
0
20
40
60
80
100
20
40
60
80
100

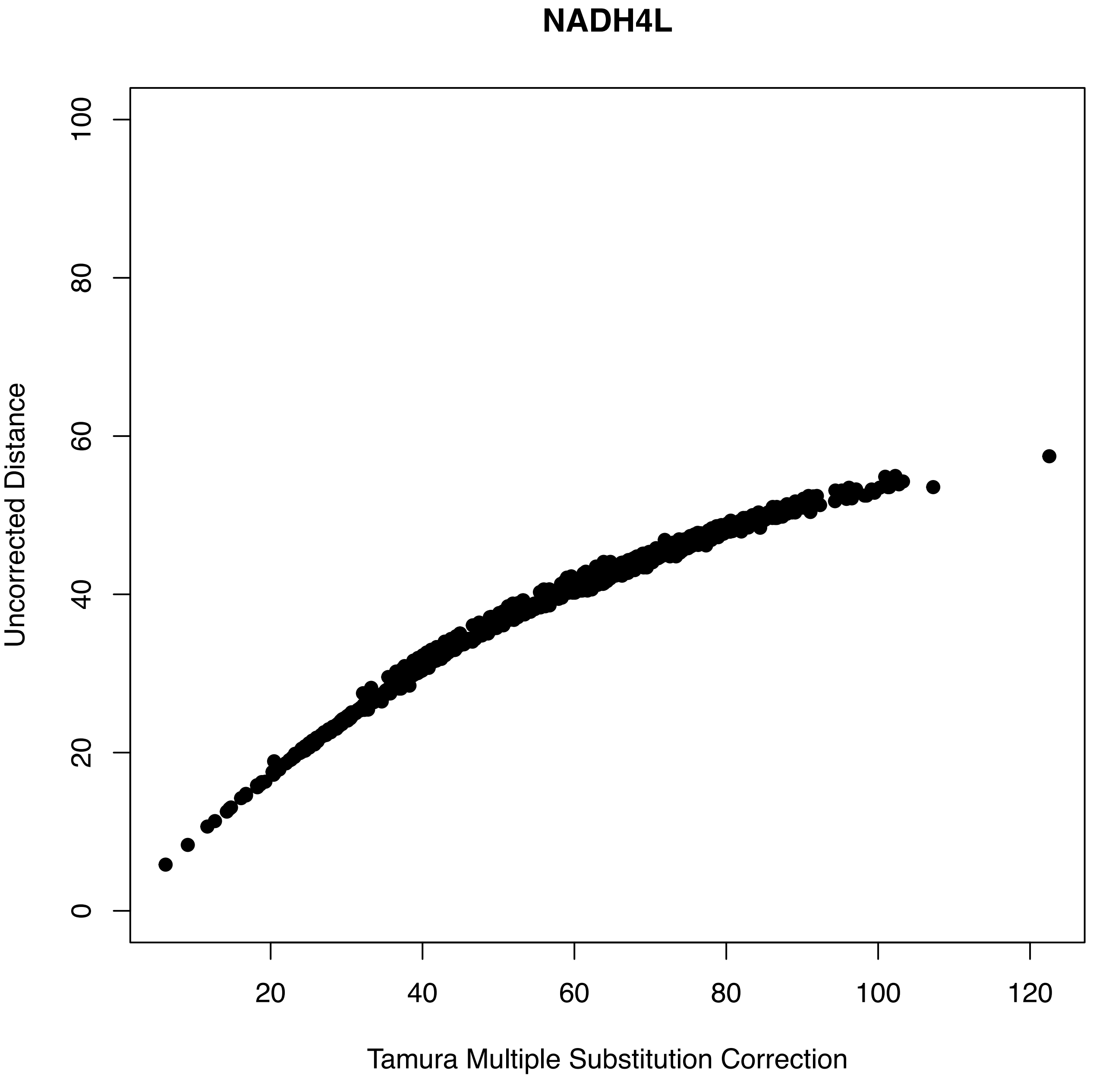
NADH4L
Uncorrected Distance
0
20
40
60
80
100
20
40
60
80
100
120
Tamura Multiple Substitution Correction

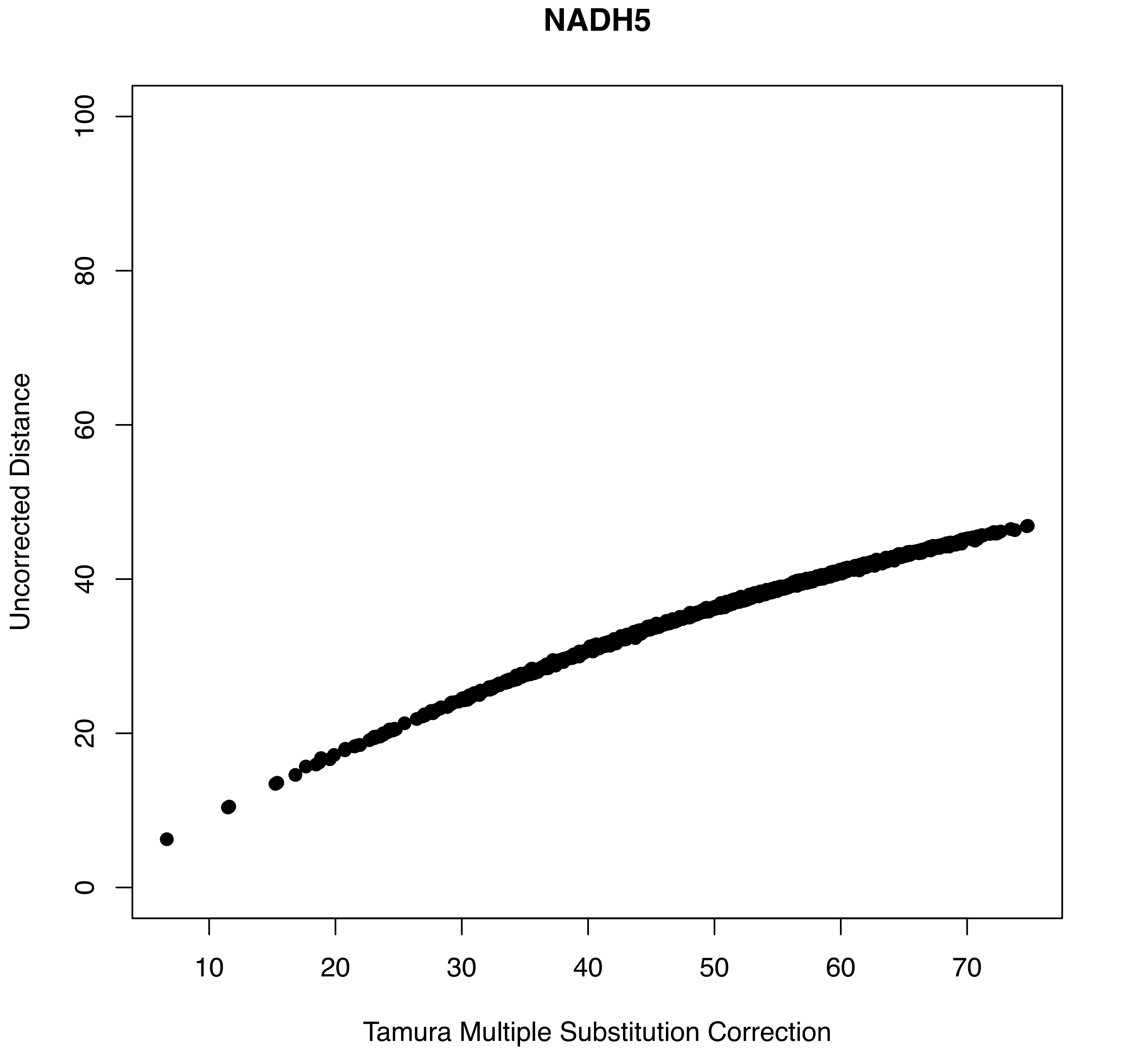
NADH5
Uncorrected Distance
Tamura Multiple Substitution Correction
0
20
40
60
80
100
10
20
30
40
50
60
70

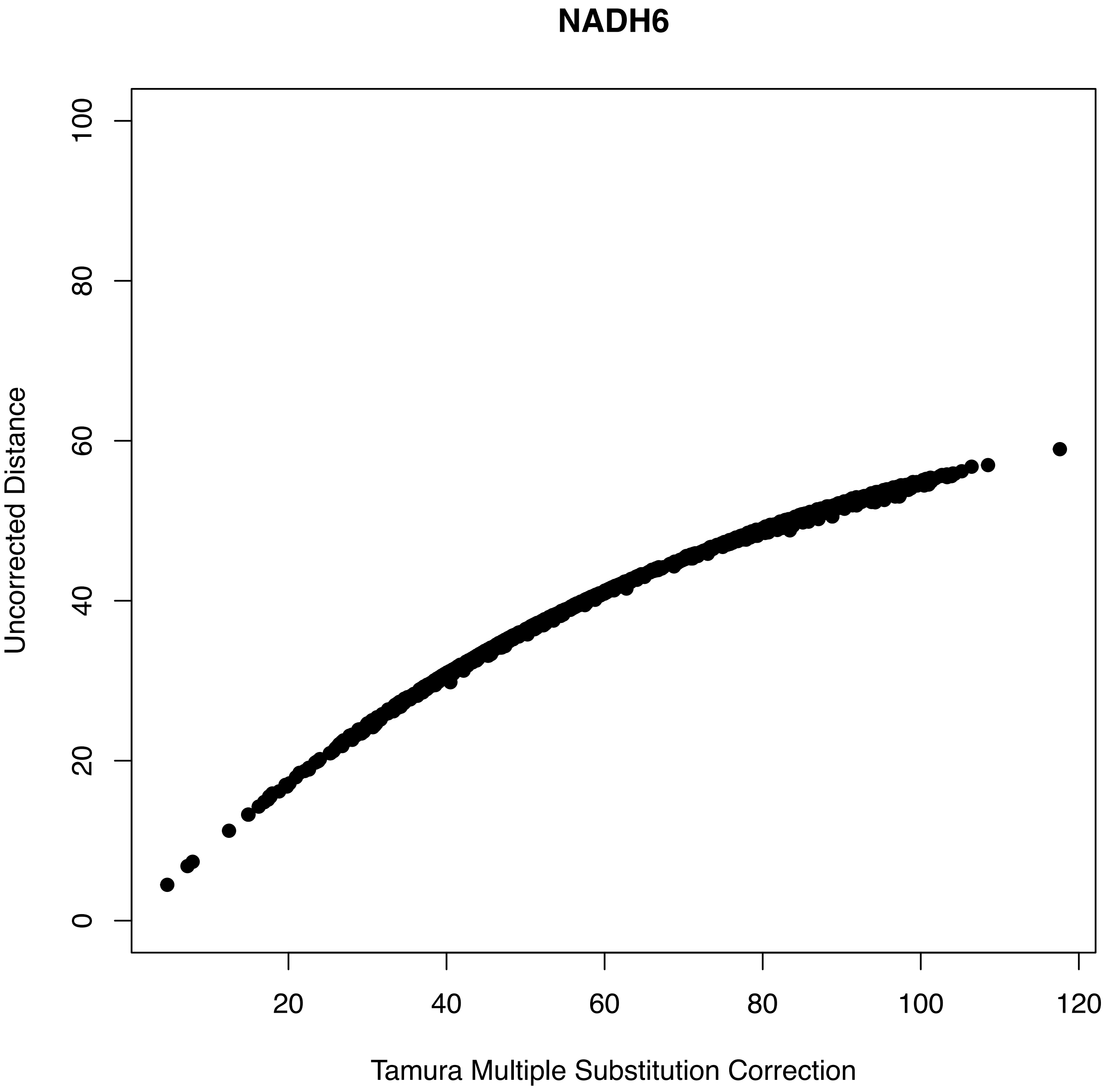
NADH6
Uncorrected Distance
0
20
40
60
80
100
20
40
60
80
100
120
Tamura Multiple Substitution Correction
