## Supplementary material for "Evidence of convergent evolution in the nuclear and mitochondrial OXPHOS subunits across the deep lineages of Squamata": S5_SSLS.pdf

**SUPPLEMENTARY MATERIALS**

**Figure S5.** Single Site Likelihood Support analysis applied on nucOXPHOS genes.
**a)** Comparisons between distributions of the first two codon positions (labeled "1") and the third codon positions (labeled "2") when ΔSSLS > 0.5. The first two codon positions show significantly higher support to Agamidae+Serpentes than the third positions (Wilcoxon rank sum test, p-value = 0.0065).
**b)** When ΔSSLS < - 0.5, comparisons between distributions of the first two codon positions ("1") and the third codon positions ("2") show no significant differences (Wilcoxon rank sum test, p-value = 0.2663).
**c)** Comparisons of ΔSSLS between contact and non-contact nucOXPHOS genes when support for one of the two topology is significant. The two groups differ significantly (Wilcoxon rank sum test, p-value = 0.000357).
**d)** ΔSSLS divided by complex. Single sites that strongly support Agamidae+Serpentes (ΔSSLS < 0.5) or Agamidae+Pleurodonta (ΔSSLS < -0.5) are highlighted in red.

**a)**

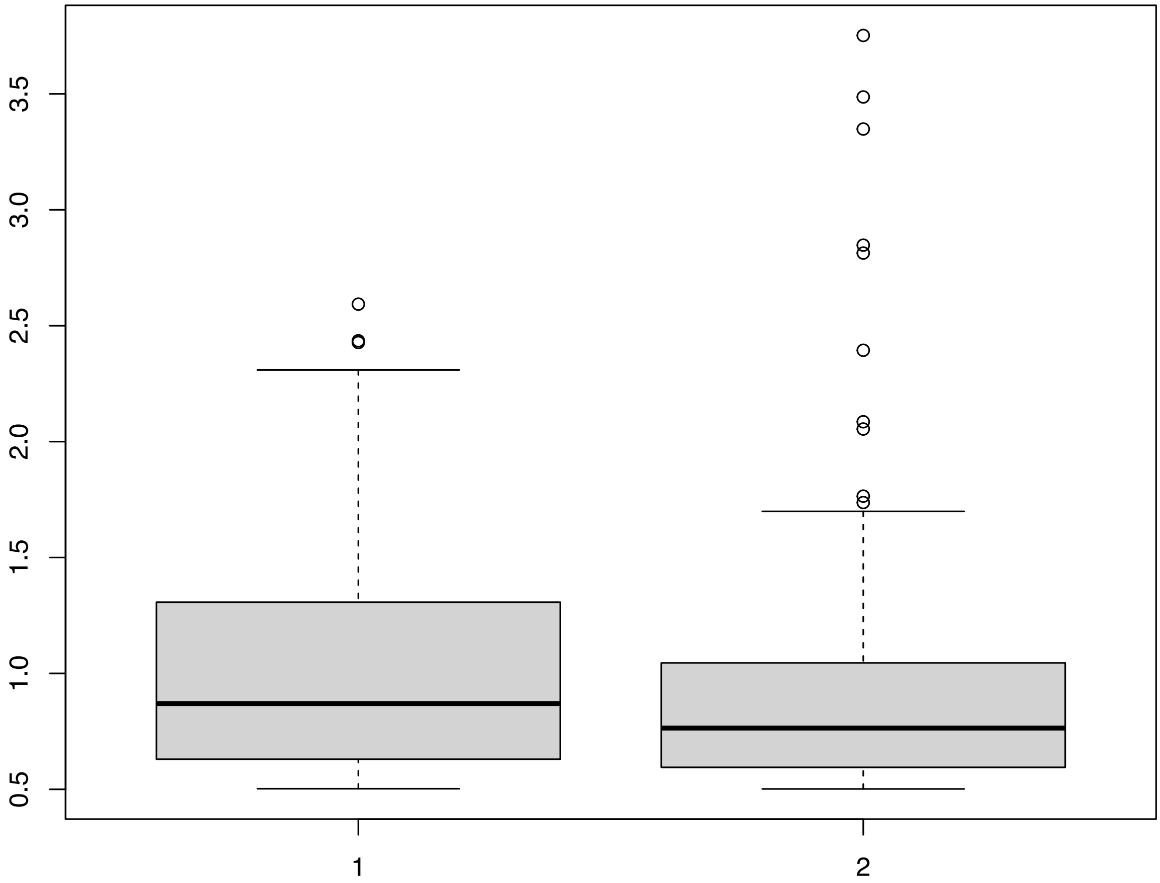

**b)**

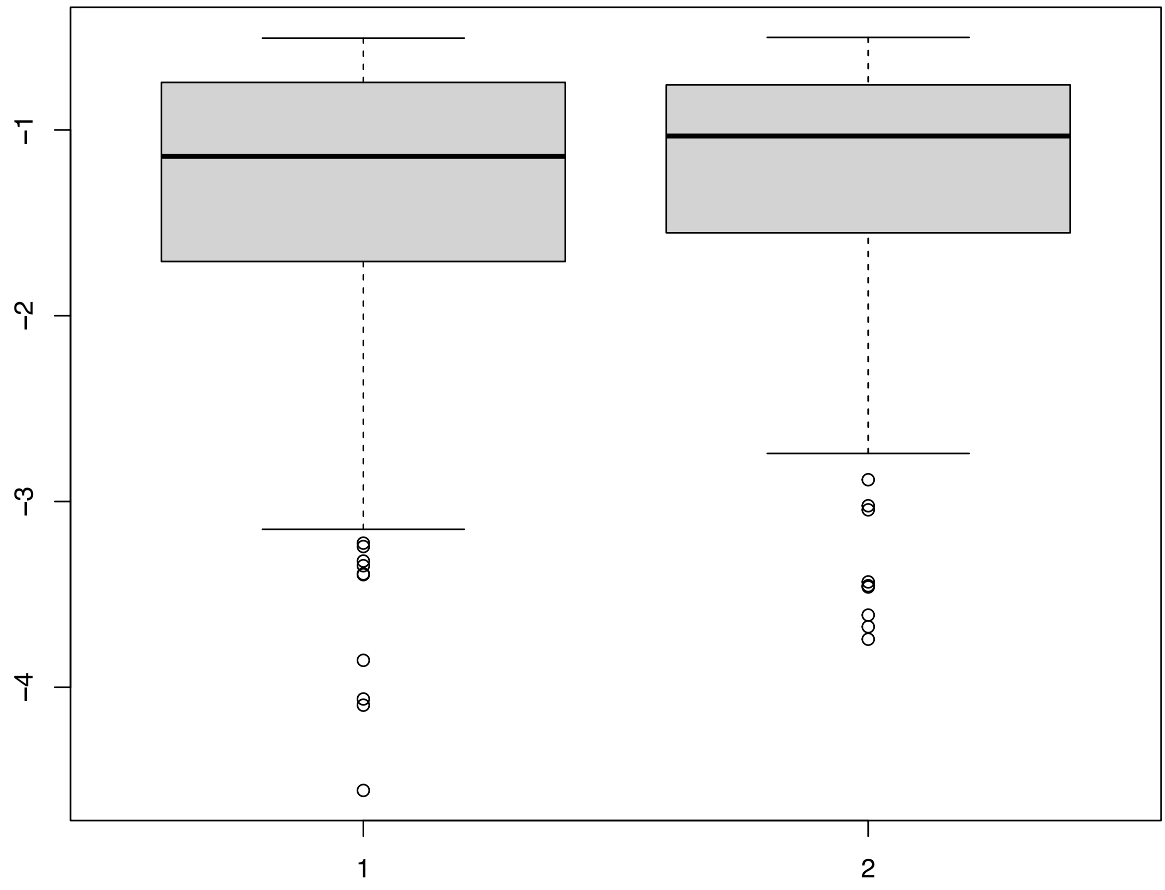


**c)**

| Type | Mean | Median | St. Dev. | N |
|---|---|---|---|---|
| Contact | -0.232 | -0.537 | 1.30 | 586 |
| Non Contact | -0.489 | -0.734 | 1.24 | 584 |

**d)**

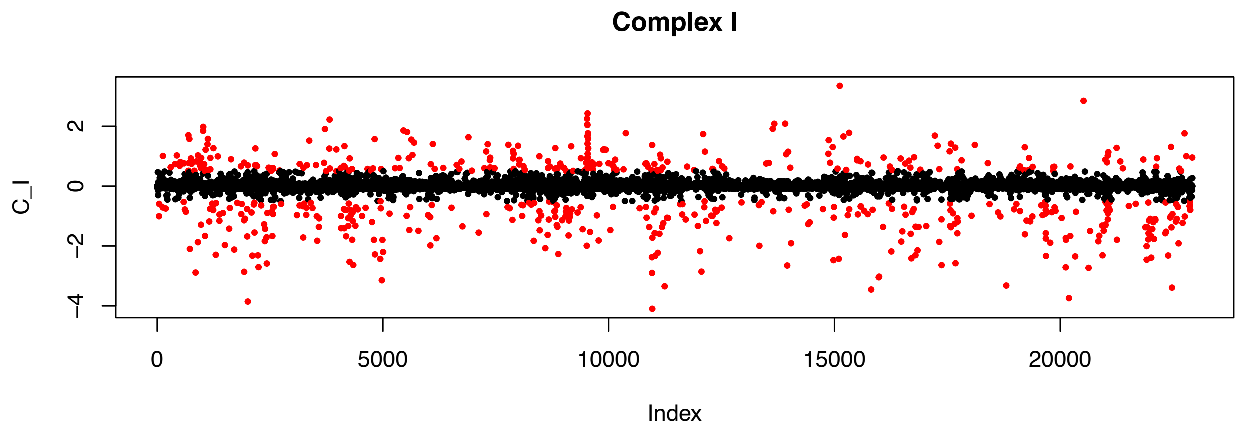

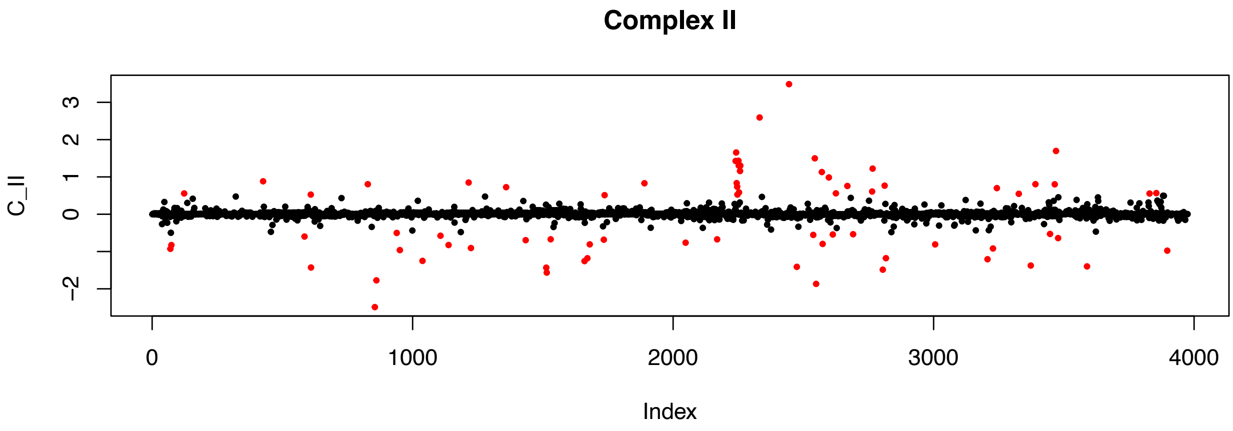
Complex II
C_II
3
2
1
0
−2
0
1000
2000
3000
4000
Index

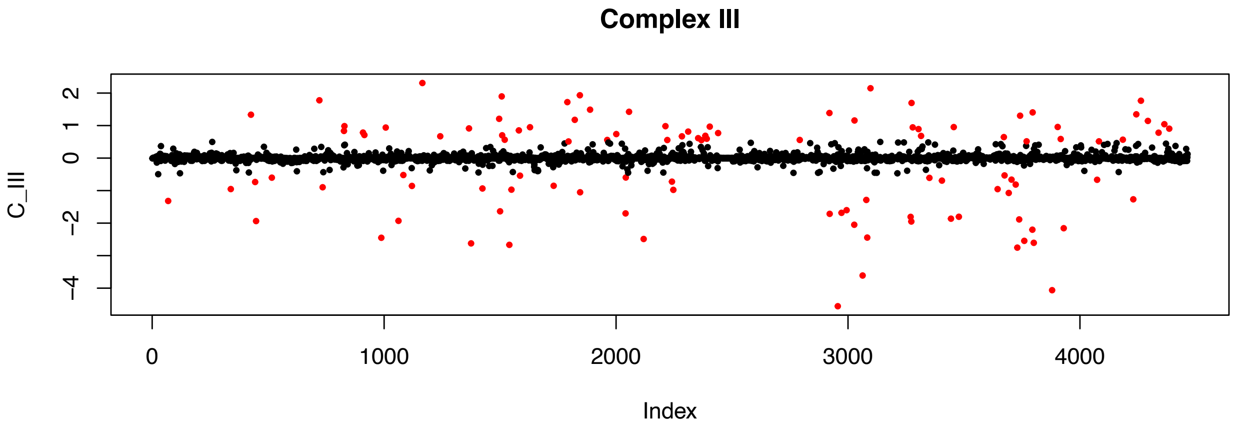
Complex III
C_III
2
1
0
−2
−4
0
1000
2000
3000
4000
Index

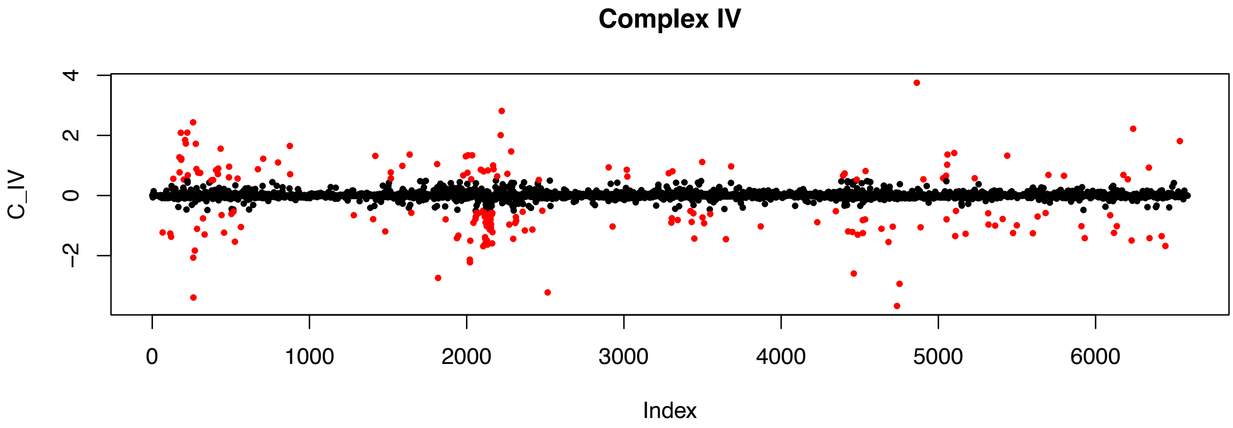
Complex IV
C_IV
4
2
0
−2
0
1000
2000
3000
4000
5000
6000
Index

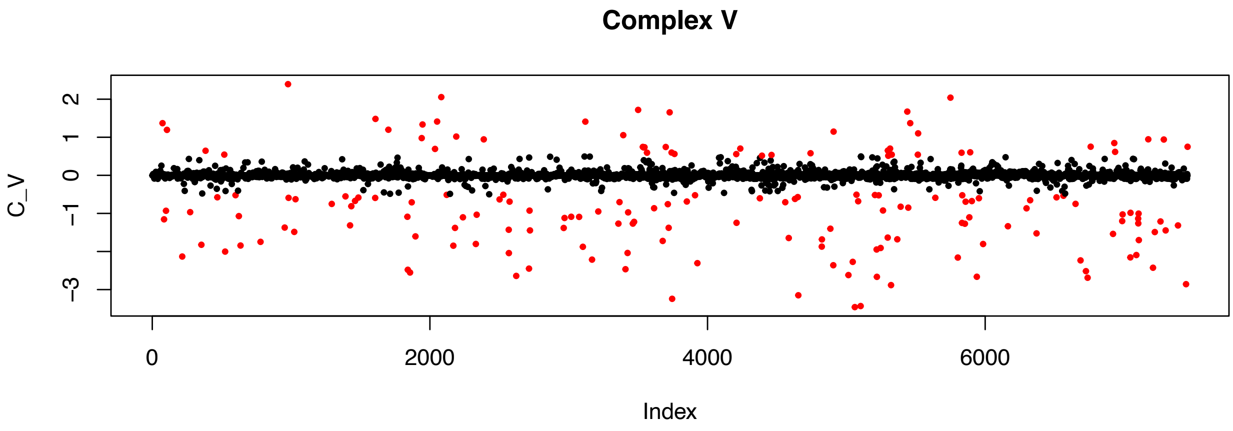
Complex V
C_V
2
1
0
−1
−3
0
2000
4000
6000
Index
