## Supplementary material for "Evidence of convergent evolution in the nuclear and mitochondrial OXPHOS subunits across the deep lineages of Squamata": S6_Flowchart_ERC.pdf

**SUPPLEMENTART MATERIALS**

**Figure S6.** Flowchart followed to perform ERC analysis, from the random extractions to the correlation plot. Visit 05_ERC for scripts.

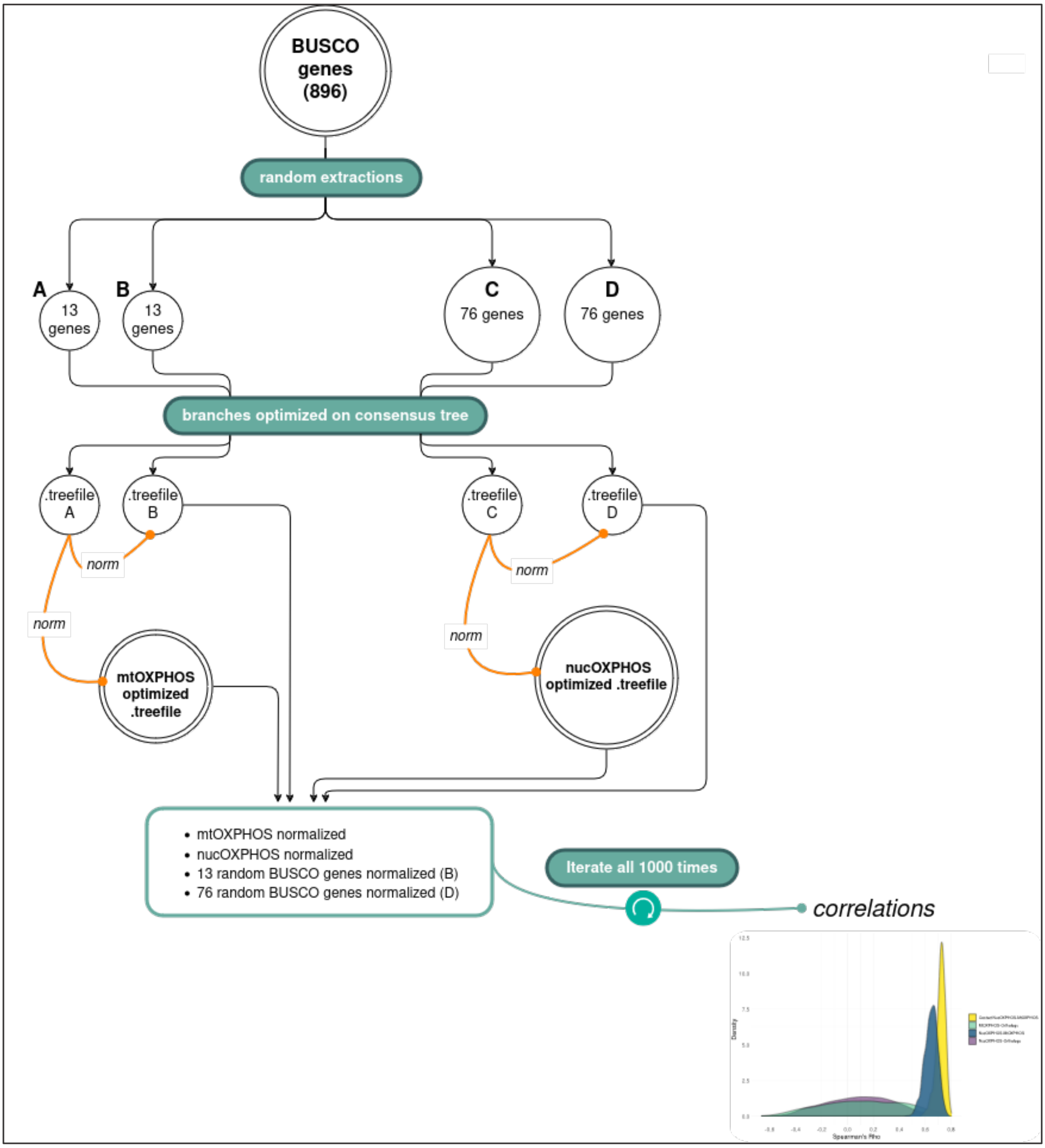
