## Supplementary material for "Evidence of convergent evolution in the nuclear and mitochondrial OXPHOS subunits across the deep lineages of Squamata": S7_Convergences.pdf

**SUPPLEMENTARY MATERIALS**

**Figure S7.** Distributions of residuals associated with **(a)** mtOXPHOS and **(b)** nucOXPHOS residuals. Residuals are defined as the distance between the observed number of convergences out of all the amino acidic changes and their expected ratio. Residual distributions were calculated along the Squamata phylogeny for each codon in each OXPHOS genes.
Points within the 95% confidence interval are shown in grey. The branch pair Agamidae-Serpentes is highlighted in red, while other colors indicate control pairs.

**a)**
mtOXPHOS

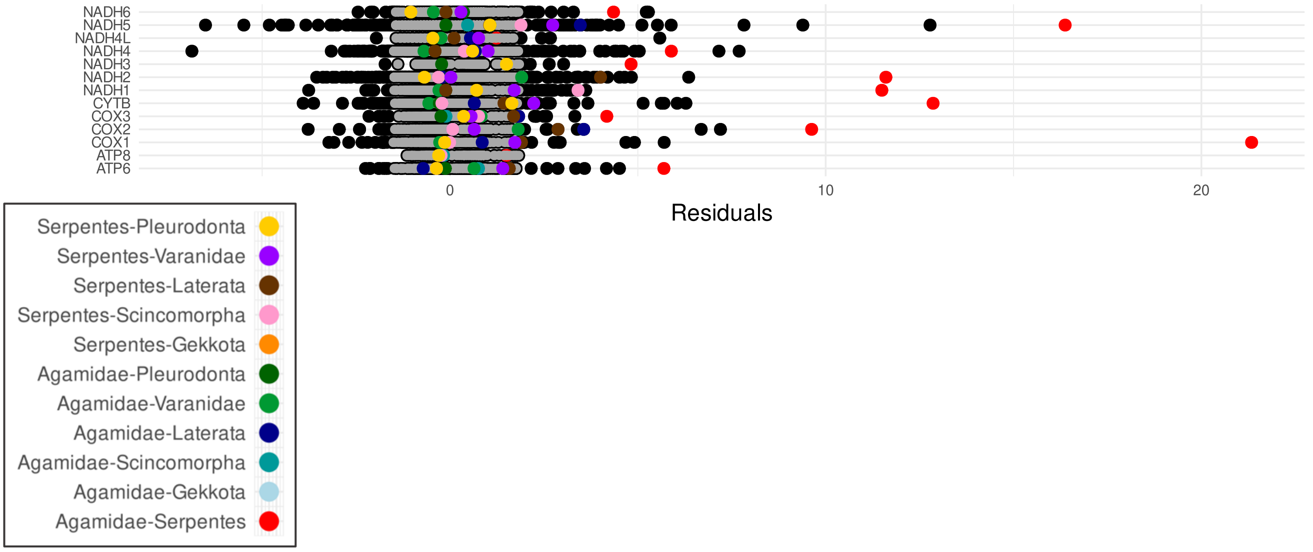

**b)**

nucOXPHOS

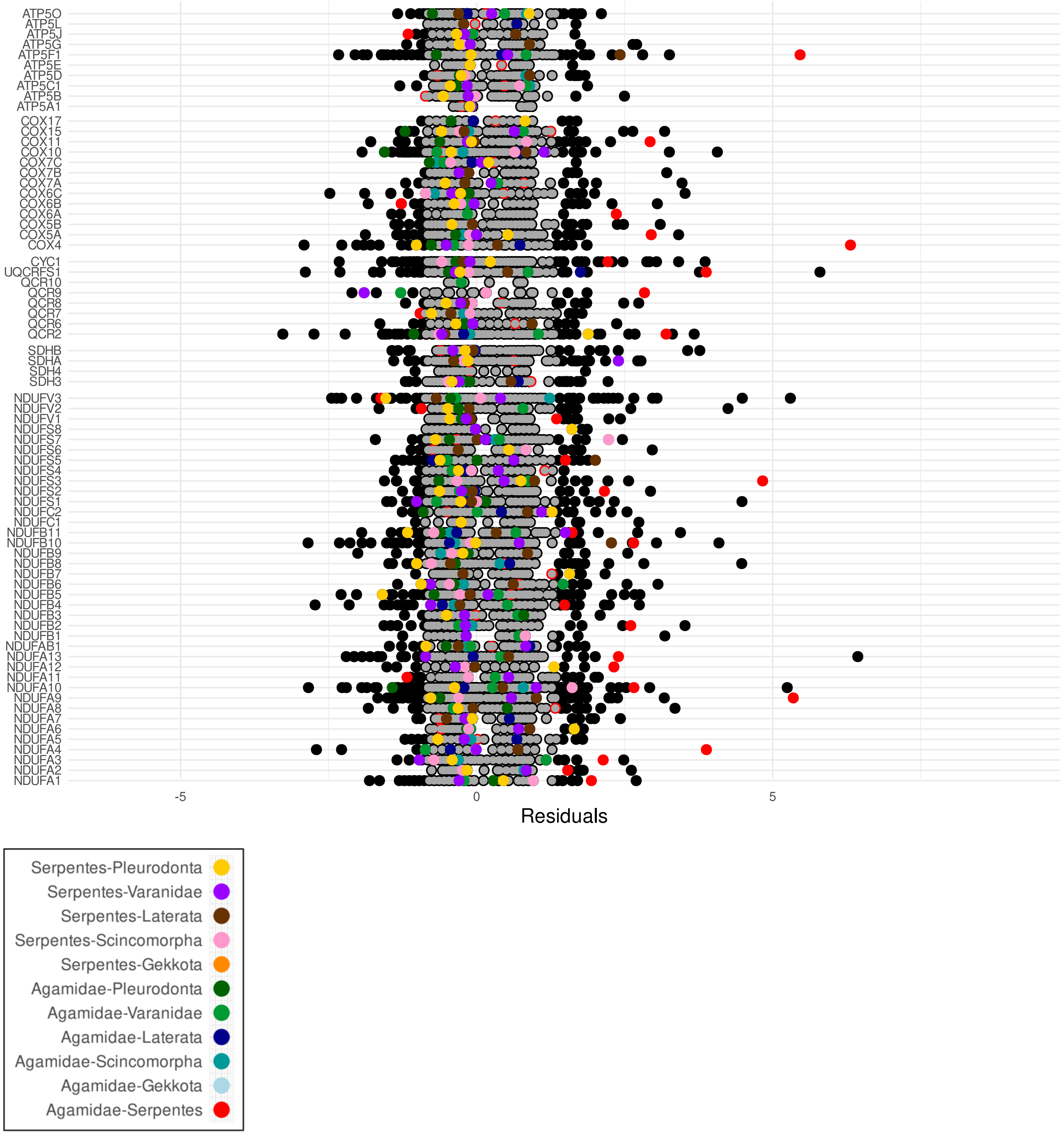

ATP5O
ATP5L
ATP5J
ATP5G
ATP5F1
ATP5E
ATP5D
ATP5C1
ATP5B
ATP5A1
COX17
COX15
COX11
COX10
COX7C
COX7B
COX7A
COX6C
COX6B
COX6A
COX5B
COX5A
COX4
CYC1
UQCRFS1
QCR10
QCR9
QCR8
QCR7
QCR6
QCR2
SDHB
SDHA
SDH4
SDH3
NDUFV3
NDUFV2
NDUFV1
NDUFS8
NDUFS7
NDUFS6
NDUFS5
NDUFS4
NDUFS3
NDUFS2
NDUFS1
NDUFC2
NDUFC1
NDUFB11
NDUFB10
NDUFB9
NDUFB8
NDUFB7
NDUFB6
NDUFB5
NDUFB4
NDUFB3
NDUFB2
NDUFB1
NDUFAB1
NDUFA13
NDUFA12
NDUFA11
NDUFA10
NDUFA9
NDUFA8
NDUFA7
NDUFA6
NDUFA5
NDUFA4
NDUFA3
NDUFA2
NDUFA1
-5
0
5
Residuals
Serpentes-Pleurodonta
Serpentes-Varanidae
Serpentes-Laterata
Serpentes-Scincomorpha
Serpentes-Gekkota
Agamidae-Pleurodonta
Agamidae-Varanidae
Agamidae-Laterata
Agamidae-Scincomorpha
Agamidae-Gekkota
Agamidae-Serpentes
