## Supplementary figures and images for "Evidence of convergent evolution in the nuclear and mitochondrial OXPHOS subunits across the deep lineages of Squamata"

### S2_Completeness_Matrices.pdf

**SUPPLEMENTARY MATERIALS**

**Figure S2.** Completeness matrices.

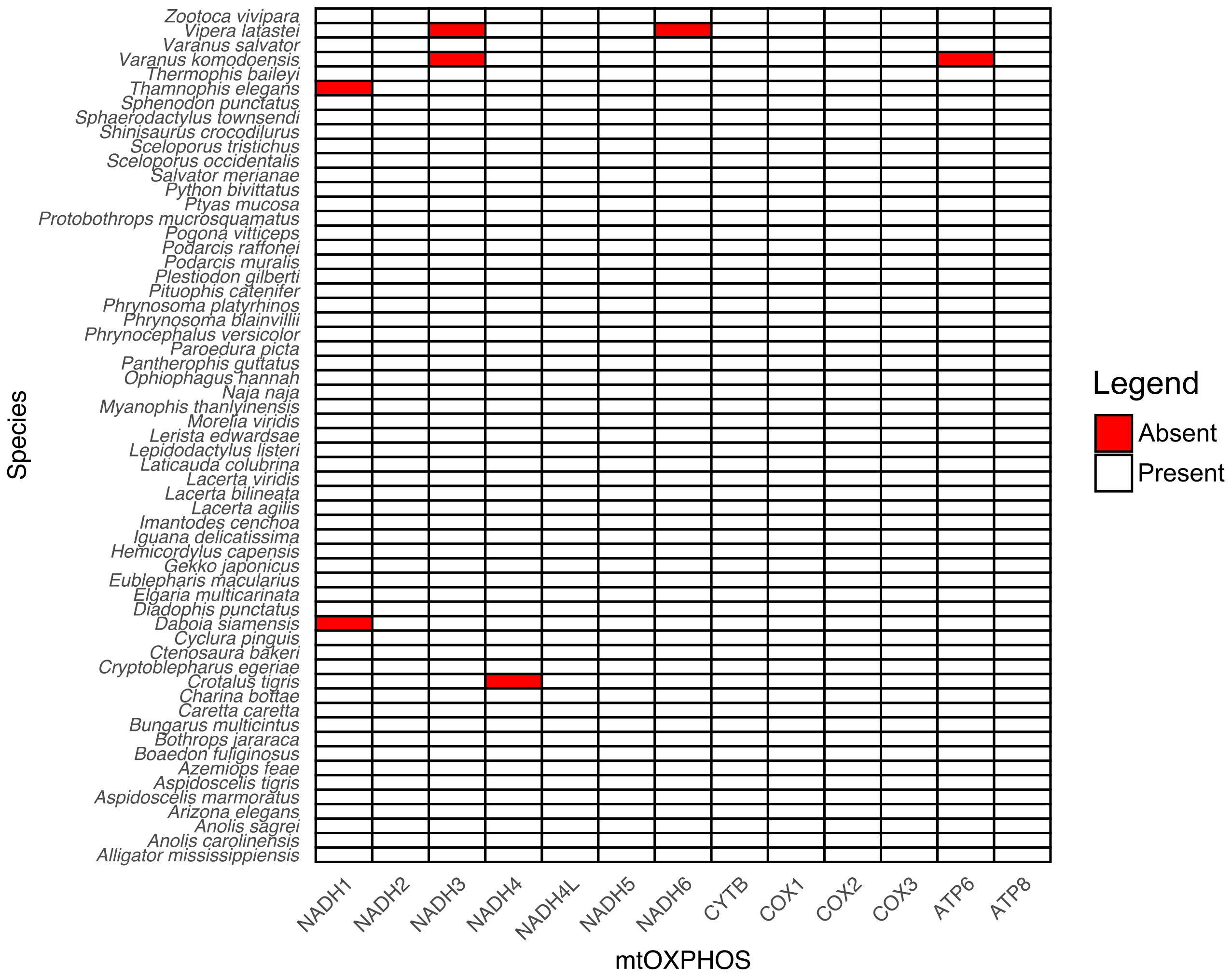

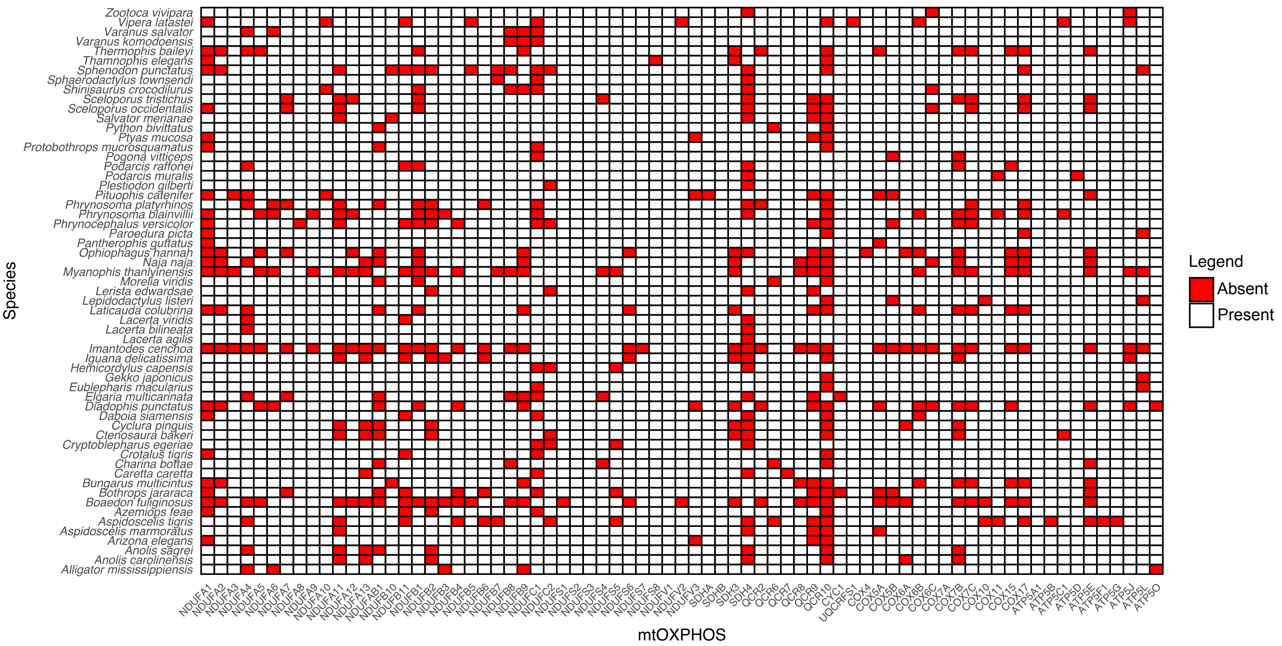
